# An efficient and robust ABC approach to infer the rate and strength of adaptation

**DOI:** 10.1101/2023.08.29.555322

**Authors:** Jesús Murga-Moreno, Sònia Casillas, Antonio Barbadilla, Lawrence Uricchio, David Enard

## Abstract

Inferring the effects of positive selection on genomes remains a critical step in characterizing the ultimate and proximate causes of adaptation across species, and quantifying positive selection remains a challenge due to the confounding effects of many other evolutionary processes. Robust and efficient approaches for adaptation inference could help characterize the rate and strength of adaptation in non-model species for which demographic history, mutational processes, and recombination patterns are not currently well-described. Here, we introduce an efficient and user-friendly extension of the McDonald-Kreitman test (ABC-MK) for quantifying long-term protein adaptation in specific lineages of interest. We characterize the performance of our approach with forward simulations and find that it is robust to many demographic perturbations and positive selection configurations, demonstrating its suitability for applications to non-model genomes. We apply ABC-MK to the human proteome and a set of known Virus Interacting Proteins (VIPs) to test the long-term adaptation in genes interacting with viruses. We find substantially stronger signatures of positive selection on RNA-VIPs than DNA-VIPs, suggesting that RNA viruses may be an important driver of human adaptation over deep evolutionary time scales.

## Introduction

Genomes contain a record of the evolutionary processes that shape diversity within and across species, and software tools that use genomic sequences to infer aspects of the evolutionary past are now an integral part of population genetics research. Of particular interest to evolutionary biologists are methods that can disentangle various processes that may contribute to diversification between species, such as adaptation and genetic drift. Such methods have the potential to resolve fundamental questions about the evolutionary (e.g. Corbett-Detig *et al*. (2015); Galtier (2016); Galtier and Rousselle (2020)) and biological (e.g. Enard *et al*. (2016); James *et al*. (2016)) drivers of diversification at the genomic level. Though numerous methods have been proposed to this end, it remains challenging to generate accurate and unbiased estimates. Studies addressing the potential biases of the available approaches unaccounted-for evolutionary processes and assessing evidence for genome adaptation is still a highly active area of research in molecular population genetics (McDonald and Kreitman 1991; Gillespie 1994; Smith and Eyre-Walker 2002; Hahn 2008; Fay 2011; Tataru et al. 2017; Kern and Hahn 2018; Jensen et al. 2019; Johri et al. 2020, 2022a,b).

The development of methods that are both computationally efficient and reasonably robust to model misspecification remains a major challenge. Most computational approaches that infer the rate of long-term adaptation at the DNA level derive from the McDonald and Kreitman (MK-test) framework (McDonald and Kreitman 1991) or the related Poisson Random Field (PRF) framework (Sawyer and Hartl 1992). Both methods use divergence and polymorphism data to estimate the proportion of non-synonymous substitutions fixed by positive selection in coding sequences, comparing alleles that are likely to have fitness effects (putatively selected) to those less likely to be under selection (putatively neutral). A significant excess of fixed differences among the putatively functional set relative to the putatively neutral set is taken as a signal of positive selection. The rate of adaptation is often summarized by the quantity *α*, which is defined as the proportion of non-synonymous (or putatively functional) fixed differences that were under positive selection along a particular evolutionary branch. Smith and Eyre-Walker (2002) applied a simple theoretical model of directional selection relating polymorphism and divergence with adaptation rate, and showed that the rate of adaptation *α* could be inferred with the quantity

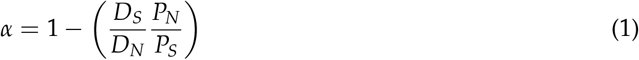

where *D*_*S*_ is the number of synonymous fixed differences in a sequencing sample, *D*_*N*_ represents non-synonymous fixed differences, *P*_*N*_ is the number of non-synonymous polymorphic sites, and *P*_*S*_ represents polymorphic synonymous sites. When *α* is close to 1, then positive selection is the predominant determinant of molecular divergence. If *α* is close to 0, then drift dominates sequence divergence. This convenient formula has been widely applied to estimate molecular adaptation, in part because of its simplicity. Indeed, the quantities on the right hand side of equation (1) are commonly inferred by comparing a population sample of sequenced individuals (sequencing sample) to a closely related outgroup species. Although widely used, it should be noted that MK-test and PRF-based approaches have multiple drawbacks that could bias the estimation of *α*. For instance, equation (1) relies on a null model derived from nearly-neutral theory (Kimura 1968; Ohta 1974; Kimura 1977), and assumes that selected polymorphism, either deleterious or beneficial, is rarely observed. Subsequent modeling and empirical studies argued that weakly selected alleles can attain both low and high frequencies and may cause substantial biases in inferences that use equation (1) (Balloux and Lehmann 2012; Lanfear et al. 2014; Booker and Keightley 2018; Galtier and Rousselle 2020; Rousselle et al. 2020). Though weakly deleterious alleles are less likely to reach fixation, they can reach relatively high frequencies and contribute to the class *P*_*N*_. This causes overestimation of the neutral mutation rate in the putatively functional category, which makes the estimation of *α* downwardly biased (Charlesworth and Eyre-Walker 2008). Fixation of weakly deleterious alleles could also cause overestimation of *α* after a long population bottleneck since the reduced population size would make the selection less efficient to purge deleterious alleles (Eyre-Walker 2002; Eyre-Walker and Keightley 2009). Conversely, weakly beneficial alleles can be found at intermediate or high frequencies (Tataru et al. 2017; Uricchio et al. 2019), especially when the rate of strongly beneficial mutations is high (as might be expected in a large population) or weakly beneficial alleles contribute substantially to polymorphism (as might be expected under some polygenic selection models). The presence of weakly selected alleles has been addressed over the last decade by MK-test and PRF-based methods mainly by explicitly modelling the Distribution of Fitness Effects (DFE) for negatively selected variants (Boyko et al. 2008; Eyre-Walker and Keightley 2009; Messer and Petrov 2013; Racimo and Schraiber 2014; Galtier 2016), along with the recent incorporation of positively selected variants in such DFE models (Galtier 2016; Tataru et al. 2017). Despite the development of several methods that account for weakly selected polymorphism, some empirical observations remain challenging to explain under existing models, such as the apparent low rate of adaptation in primates, the constrained range of genetic diversity across species, and differences in the rate of adaptation among taxa (Galtier 2016; Castellano et al. 2018, 2019a). Generating a deeper biological and evolutionary understanding of the drivers of differentiation across species may require new methods and models that can efficiently estimate the DFE while simultaneously accounting for many (potentially confounding) evolutionary processes.

Demographic processes (such as population contractions, expansions, and migrations) are another primary potential source of bias in the inference of selection (Jensen et al. 2019; Johri et al. 2020, 2022a,b), just as the selection is an essential potential confounder in the inference of demography (Schrider et al. 2016; Torres et al. 2018). The developers of robust inference methods have typically sought to account for both selection and demographic processes simultaneously. The cost of incorporating both demography and selection is accrued in terms of model complexity and loss of efficiency, as it is much more challenging to compute likelihoods or summary statistics under joint demography/selection models. There is some hope, however, that methods based on the asymptotic MK-test (aMK-test) (Messer and Petrov 2013) may have some inherent robustness since these approaches rely on summary statistics that involve ratios of functional and (putatively) non-functional alleles. Hence, some of the effects of demography should be absorbed into the ratio, as both categories of alleles will be affected. Here, we develop an extension of the Approximate Bayesian Computation ABC-MK method presented in Uricchio *et al*. (2019) that greatly improves the efficiency of the population genetics inferences. In Uricchio *et al*. (2019), analytical calculations were used to explore the effect of background selection (BGS) and selective interference on weakly beneficial alleles, but the estimation procedure employed was based on computationally intensive forward-in-time simulations and took days even on a High Performance Computing cluster. We developed a simpler and much more computationally efficient ABC-based inference procedure that accounts for the DFE of deleterious and beneficial alleles and partial recombination between selected genomic elements while avoiding forward-in-time simulations. We describe the inference procedure, assess its performance and robustness to non-equilibrium demographic scenarios and different intensities of adaptation, and apply it to human genomic data. We show that the method is reasonably robust to non-equilibrium events or different fitness values of adaptation, use it to provide additional evidence for a substantial effect of different RNA-viruses on human adaptation rates, and discuss caveats and potential extensions of our work.

## Methods

### Preliminaries

Our first goal is to calculate the expected rate of fixation and the expected site frequency spectrum of neutral and selected polymorphism under a model of directional selection with incomplete recombination. To do so, we follow the results of Uricchio *et al*. (2019), which in turn extended the results of several earlier studies (e.g., Eyre-Walker and Keightley (2009); Messer and Petrov (2013)). In subsequent sections we extend these calculations by developing a random sampling scheme that accounts for the Poisson variance in mutation and fixation rates, and allows us to develop a simple inference pipeline. We briefly review the core aspects of the theoretical framework.

Our ultimate goal is to estimate *α*, the proportion of non-synonymous substitutions fixed by positive selection, as well as the DFE over de novo non-synonymous mutations and substitutions. We suppose that selection is directional, with both positively selected and negatively selected mutations. We first consider the case where each selected locus evolves independently, and in subsequent sections we consider cases with BGS and selective interference.

The DFE over beneficial alleles consists of two point masses, one representing strongly beneficial alleles and the other representing weakly beneficial alleles. The rate of adaptation can therefore be decomposed into weakly and strongly beneficial components, *α* = *α*_*W*_ + *α*_*S*_. The substitution rate for non-synonymous alleles is denoted as *d*_*N*_, with 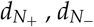, and 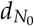representing the rates for positively selected, negatively selected, and neutral alleles respectively (note that 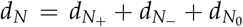 by definition - since there is a DFE for beneficial and deleterious alleles we could further decompose 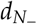 and 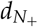 into more categories, but we collapse the DFE into two compartments here and consider the full DFE in later sections). In the same way, we denote as *d*_*S*_ the substitution rate of synonymous mutations, which are assumed to be neutral. We can write *α* as

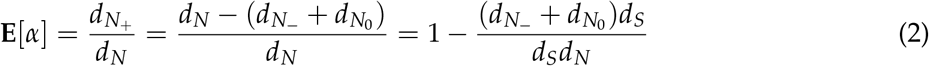

Note that we define *α* as the realized proportion of positively selected substitutions along the branch, and hence equation (2) is an expression for the expectation of *α*. As noted by Messer and Petrov (2013), *d*_*S*_ can be estimated from sequence alignments with the ratio *D*_*N*_/*D*_*S*_ under the assumption that the observed number of substitutions along a branch should be proportional to the rate. The ratio 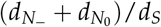 is more complex to estimate, because it relies on partitioning substitutions by their fitness effects. Under the assumption that polymorphic alleles are rarely selected (because deleterious sites are removed from the population quickly and beneficial sites go to fixation rapidly), previous work (Smith and Eyre-Walker 2002) showed that this ratio can be approximated by substituting 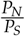 into equation (2), and a point estimate of *α* as

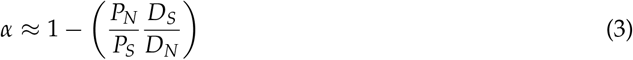

However, if selected polymorphisms segregate in the sample, then *P*_*N*_ in equation (3) will be inflated relative to the true rate of mutation for neutral non-synonymous alleles, which results in underestimation of *α*. A potential solution is to exclude alleles with derived allele frequencies lower than some threshold from the quantities *P*_*N*_ and *P*_*S*_, since most (negatively) selected alleles should be constrained to lower frequency (Fay et al. 2001). While this solution works well for some DFEs (for example, when all deleterious alleles are strongly selected), weakly deleterious alleles can reach appreciable frequencies and bias inference regardless of the selected frequency threshold. Messer and Petrov (2013) extended this idea by developing a very simple estimator of *α* (asymptotic-MK) that uses all frequencies simultaneously by rewriting the estimator of Smith and Eyre-Walker (2002) as

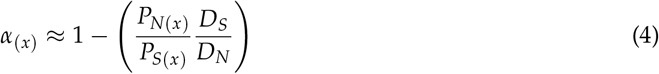

where *P*_*N*(*x*)_ and *P*_*S*(*x*)_ are the number of non/synonymous alleles at frequency *x* in a sequencing sample. This method improves the quality of *α* estimates by using all of the frequency data simultaneously and providing confidence intervals for *α*. The estimate of *α* results in an exponential function of the form: *α* = *a* + *b · e*^−*cx*^, where the best fit of the exponential at *x* = 1, eliminates the effect of a weakly deleterious allele. In the following section, including empirical and analytical results, we defined *P*_*N*(*x*)_ and *P*_*S*(*x*)_ as all non-synonymous and synonymous polymorphisms above frequency *x*. As noted in Uricchio *et al*. (2019) this approach scales better as sample size increases since most common allele frequencies *x* have very few polymorphic sites in large samples and both *P*_*N*(*x*)_ and *P*_*S*(*x*)_ quantities trivially have the same asymptote. Despite the improvement, the method does not estimate the DFE and assumes that beneficial alleles do not contribute to *P*_*N*_, which turns into *α* underestimation in cases where selection is predominantly weak.

### Expected fixation rates and frequency spectra

A complementary approach to that of Messer and Petrov (2013) is to directly model the effects of the DFE for beneficial and deleterious alleles on the shape of the *α*_(*x*)_ curve, and to infer the best fitting model parameters. Nonetheless, note that while 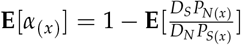 is not straightforward to calculate because it depends on the ratio of several random variables, the expectation of each component in equation 4 (*P*_*S*(*x*)_, *P*_*N*(*x*)_, *D*_*S*_, *D*_*N*_) is easily calculated in a directional selection model from first principles using diffusion theory (Evans et al. 2007). Therefore, we make a first-order approximation

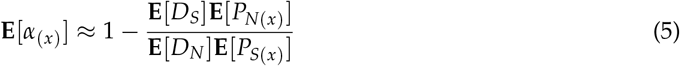

Our model will assume that positively selected mutations have fitness effects drawn from a point mass distribution with two values defining either strong or weak adaptation (although such an assumption is relaxed in the Approximate Bayesian Computation), and negatively selected mutations were Gamma-distributed following previous studies (Eyre-Walker et al. 2006; Boyko et al. 2008; Eyre-Walker 2010), replacing *θ*_*s*_ = 4*Nµs* with Gamma distribution Γ[*α, β*] over selection coefficients in equation (6) and (7).

In general, our approach can be applied to any distribution for which we can analytically solve the fixation rates and expected frequency spectra. Descriptions of these calculations follow this section and can be found in the online documentation for our software at web address jmurga.github.io/MKtest.jl.

### Expected frequency spectrum

The expected number of alleles at frequency *x* is estimated from the standard diffusion theory for the site frequency spectrum in an equilibrium population (e.g., see equation 31 of Evans *et al*. (2007)).

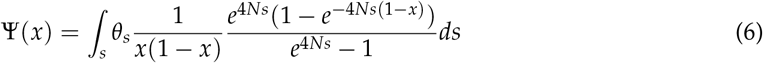

where Ψ(*x*) is the expected alleles at frequency *x* in a population of size *N* and *θ*_*s*_ = 4*Nµs* is the population-scaled mutation rate for mutations with selection coefficient *s*. To obtain the downsampled frequency spectrum in a finite sample of 2*n* chromosomes, we convoluted equation (6) with the binomial distribution.

### Expected number of fixations

Considering the distribution of selection coefficients over new mutations *µ*_*s*_ (selection coefficient underlying the mutation rate) and the fixation probability *π*_*s*_, we calculate the expected fixation rate as

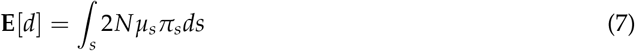

Following Uricchio *et al*. (2019), the expected fixation rate can be decomposed by its fitness effect replacing *θ*_*s*_ = 4*Nµs* in equations (6) and (7) with a Gamma distribution Γ[*α, β*] over negative selection coefficients and point mass distribution over positive selection coefficients,

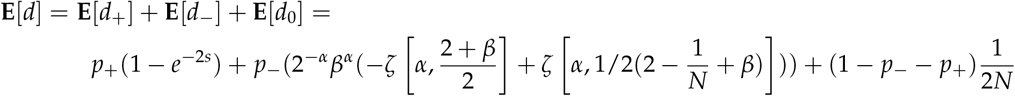

where *p*_+_ and *p*_−_ are the probability that an allele is beneficial or deleterious, respectively, and *ζ* is the Riemann Zeta function (Eyre-Walker 2010).

### Background selection and adaptive divergence

BGS (Charlesworth et al. 1993; Hudson and Kaplan 1995; Nordborg et al. 1996) and selective interference (e.g., Hill-Robertson interference, (Hill and Robertson 1966)) could affect the rate of fixation of weakly deleterious or beneficial alleles. Up to this point, we have considered only selected loci that evolve independently of all other selected loci. In this section we will relax this assumption by exploring the effects of selective interference on fixation rates and the frequency spectrum. We will follow approximations that will apply in some circumstances (in particular, when BGS is the predominant driver of selective interference), but may fail when strongly beneficial alleles interfere (Good et al. 2014; Cvijovic’ et al. 2018; Garcia and Lohmueller 2021; Ragsdale 2022).

To explore *α*(*x*) accounting for recombination and BGS impact, we focused on a model in which the coding locus is flanked on each side by loci of length *L*, which contain deleterious alleles. We modeled deleterious alleles with a population-scaled selection coefficient −2*Nt* undergoing persistent deleterious mutation at rate 4*Nµ*_−_ and the whole flanking loci recombined at a rate *r* per-base, per-generation. The effects of BGS on fixations and frequency spectra have been subject of much theoretical work (Charlesworth et al. 1993; Charlesworth 1994; Hudson and Kaplan 1995; Barton 1995; Nordborg et al. 1996). Previous work has shown that diversity at the coding locus (*π*) is decreased relative to its neutral expectation (*π*_0_), and closed form expressions for the expected reduction in diversity are available (Hudson and Kaplan 1995; Nordborg et al. 1996). While patterns of sequence variation induced by BGS can be quite complex (Nicolaisen and Desai 2013; Good et al. 2014; Torres et al. 2018, 2020), to a first approximation the effect of BGS can be thought of as a reduction in the effective population size *N*_*e*_, with 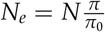 (McVicker et al. 2009). To account for the role of BGS on the fixation rate of deleterious alleles, we replace *N* in the prior equations with *N*_*e*_ after accounting for BGS. We also replace *N* with *N*_*e*_ in formulae for the frequency spectra.

For beneficial alleles, the effects of selective interference are slightly more complex. Strongly beneficial alleles are essentially unaffected by BGS, in that their fixation probabilities almost do not depend on the reduction in neutral diversity (Uricchio et al. 2019). Weakly beneficial alleles can have their fixation probabilities substantially reduced by BGS. We followed Barton (1995) to derive formulae for the reduction in fixation rate of weakly and strongly beneficial alleles after accounting for BGS, as described in the Supplemental Materials of Uricchio *et al*. (2019) (see *Background selection and adaptive divergence* section). The reduction in fixation probability for a weakly beneficial allele with selection coefficient *s* under interference with deleterious alleles with selection coefficient *t* is given by

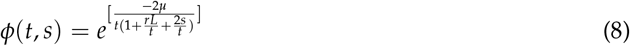

where *l* is the distance in base pairs from the region of interest, 1 ≤ *l* ≤ *L* (see equation 17d of Barton (1995)). Multiplying across all deleterious linked sites and factoring in flanking sequences to both the left and right of the focal site (which requires us to square the product below), we find that the reduction in the probability of fixation relative to the case with no linkage (Φ) is

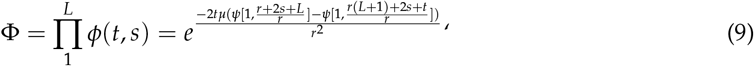

where *ψ* is the polygamma function. We note that these expressions are not expected to hold under very high rates of mutation for beneficial alleles, and will be less accurate for strongly beneficial alleles than weakly beneficial alleles. Given these expressions, we can replace the fixation rates for beneficial alleles in our prior formulae with the fixation rates after accounting for selective interference.

### Poisson-sampling process

The previous sections described the expectation of fixation rates and frequency spectra under a model of directional selection and selective interference. We now develop a simple random sampling scheme around these expectations that accounts for sampling and process variance, linking analytical calculations to an ABC procedure which avoids computationally expensive forward simulations. We note that the model we explore is quite similar to the BGS model in DeGiorgio *et al*. (2016), though while we are interested in the long-term accumulation of fixations, DeGiorgio *et al*. (2016) is primarily interested in the non-equilibrium signature of a recent or ongoing selective sweep. Following the Poisson Random Field model (Sawyer and Hartl 1992), we supposed that the number of fixed differences and polymorphic sites were Poisson random distributed variables with mean values given by the expectations in the previous sections.

To avoid performing branch length estimations in our computation, we assumed that the empirically observed number of fixations should be proportional to the length of the evolutionary branch of interest, *T*, the locus length *L* and mutation rate *µ*. We take the observed total number of fixations (including both non-synonymous and synonymous sites) as a proxy for the expected number, and then sample weakly deleterious, neutral, and beneficial substitutions proportional to their relative expected rates for a fixed set of model parameters. The expected number of substitutions for positively selected substitutions is then

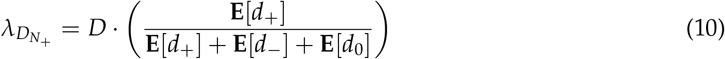

where *D* is the observed number of substitutions in a dataset of interest.

It should be noted that both sampling variance and process variance affect the number of variable alleles at any particular allele frequency in a sequencing sample. The process variance arises from the random mutation-fixation process along the branch, while the sampling variance arises from the random subset of chromosomes that are included in the sequencing data. We sampled a Poisson distributed number of polymorphic alleles at frequency *x* relative to their rate given the expected frequency spectra. The expected frequency spectra were downsampled using a binomial (with probability of success given by the frequency 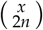 in a sample of 2*n* chromosomes) to account for the sampling variance. In a manner exactly analogous to fixed variants as described above, to account for the process variance

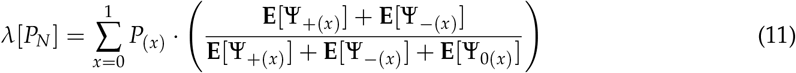

To account for background selection, we solved equation (9) using any expected *B* values from McVicker *et al*. (2009) at each polymorphic or fixed site. Then, we adjust the fixation probability of deleterious alleles by the expected value of *B* and the fixation probability for weakly beneficial alleles given the predicted reduction by equation (9). In practice, we bin sites into *B* value ranges, such that all sites within (for example) a 2.5% *B* value range of 0.675 to 0.7 experience the same *N*_*e*_ and the same strength of selective interference (for example, *N*_*e*_ = 0.675*N*, which is the midpoint of this *B* value window). Because we note that inferred background selection strength is unavailable for most species, our software can run without this information by inputting any expected *B* value range into the software. Nonetheless, the analysis can be limited to an empirical *B* value or *B* value range as an approximation of the expected reduction in diversity if downstream empirical datasets account for the inferred background selection strength estimated from McVicker *et al*. (2009) or Murphy *et al*. (2022).

### Computational workflow

Our goal is to infer *α, α*_*W*_, and *α*_*S*_ given a set of observed *α*_(*x*)_ values from a sequencing dataset. Since *α* in our framework is a model output and not a parameter per se (i.e., *α* depends on the random number of fixations along an evolutionary branch, which in turn depend on the parameters of the evolutionary model), we cannot immediately obtain the corresponding fixation rates and frequency spectra values for a given set of expected *α* values without first solving for the mutation rates and fixation probabilities considering the input model (see Table 1). Given the *α* and *α*_*W*_ (which together uniquely determine *α*_*S*_), a DFE over negatively selected alleles, a known *B* value, a selection coefficient for beneficial alleles, a selection coefficient for flanking deleterious alleles, a recombination (*ρ*) and mutation rate on the coding locus (*θ*), we numerically solve for the probability of fixation of beneficial alleles and the mutation rate on the flaking locus that correspond to the desired *α* values and BGS strength. This allows us to rapidly calculate the expected frequency spectra and fixation rates that will correspond to the desired *α* values and generate a sample of *α*_(*x*)_ values following the Poisson-sampling process under the corresponding evolutionary model.

**Table 1.** Model parameter definitions.

|  |  |
| --- | --- |
| $\gamma$ | Population-scaled selected coefficient of deleterious alleles |
| $s_w$ | Population-scaled selected coefficient of weakly beneficial alleles |
| $s_S$ | Population-scaled selection coefficient of strong beneficial alleles |
| $\alpha_W$ | Proportion of weakly adaptive substitutions |
| $\alpha$ | Proportion of adaptive substitutions |
| $\theta_{flaking}$ | Population-scaled mutation rate at flanking non-coding locus |
| $\theta_{coding}$ | Population-scaled mutation rate at coding locus |
| $\theta$ | Scale parameter |
| $\beta$ | Shape parameter |
| $B$ | $B$ value from \cite{mcvicker_widespread_2009} |
| $l_w$ | Fixation probability of weakly beneficial alleles |
| $l_S$ | Fixation probability of strong beneficial alleles |
| $N$ | Effective population size |
| $n$ | Sample size |
| $L$ | Flanking locus length |
| $t$ | Population-scaled selected coefficient of flanking deleterious alleles |

### Inferring parameters with ABC

We used a generic Approximate Bayesian Computation (ABC) algorithm to infer the rate and strength of adaptation. ABC first samples the parameter values from prior distributions; second, simulate the model using these random parameter values and calculates informative summary statistics from the simulations; and third, compares the simulated summary statistics to observed empirical data. The summary statistics that best match the observed empirical data form an approximate parameter posterior distribution. Our approach used empirical data to both perform computational workflow and the Poisson-sampling scheme described above to sample *α*_(*x*)_ generating summary statistics corresponding to different evolutionary scenarios and to finally compare such summary statistics to empirical *α*_(*x*)_ estimations. Since we do not know a priori the values of any model parameter, to estimate summary statistics, we sample 10^5^ sets of parameters randomly from a prior uniform distribution (see Table 1, Table 2), which allows for flexibility in the DFE of deleterious and beneficial alleles. We supplied the summary statistics and empirical *α*_(*x*)_ into ABCreg (Thornton 2009) to estimate the empirical values of *α*_*W*_, *α*_*S*_ and *α* while accounting for BGS in bins of 2.5% from 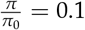to 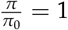.

**Table 2.** ABC-MK prior values.

| Scenarios | $N$ | $n$ | $\alpha$ | $s_W$ | $s_S$ | $\gamma$ | $\beta$ | $t$ | $B$ |
| --- | --- | --- | --- | --- | --- | --- | --- | --- | --- |
| Demographic equilibrium | 500 | 500 | [0.1,0.9] | [1,10] | [200,2000] | [-1000,-200] | $0.184 \cdot 2^{[-2,2]}$ | [-1000,-500] | [0.1,0.999] |
| Tennensen | 1000 | 661 | [0.1,0.9] | [1,10] | [200,2000] | [-1000,-200] | $0.184 \cdot 2^{[-2,2]}$ | [-1000,-500] | [0.1,0.999] |
| Twoepoch B=0.4 | 1000 | 50 | [0.1,0.5] | [1,10] | [200,2000] | [-1000,-200] | $0.184 \cdot 2^{[-2,2]}$ | [-1000,-500] | [0.3,0.5] |
| Twoepoch B=0.999 | 1000 | 50 | [0.1,0.5] | [1,10] | [200,2000] | [-1000,-200] | $0.184 \cdot 2^{[-2,2]}$ | [-1000,-500] | [0.8,0.999] |
| Thousand Genomes Project data | 1000 | 661 | [0.1,0.9] | [1,10] | [200,2000] | [-1000,-200] | $0.184 \cdot 2^{[-2,2]}$ | [-1000,-500] | [0.1,0.999] |

We conducted ABC inferences using two different kinds of data, as explained in the following sections. First, for simulated data, we bootstrapped the simulated replicas to generate 100 datasets to test the method. For empirical data, we followed the bootstrap procedure extensively described at Enard and Petrov (2020) and Di *et al*. (2021) to generate the control sets required to create null distributions and compare adaptation levels between non-VIPs and VIPs genes.

As summary statistics, we used the value of *α*_(*x*)_ for *x* ∈ (2, 5, 20, 50, 200, 661, 925), *x* ∈ (2, 5, 20, 50, 200, 500, 700), *x* ∈ (2, 4, 5, 10, 20, 30, 50, 70) regarding the input data and sample size. These values are proportionally similar to the ones used in Uricchio *et al*. (2019). However, we excluded singletons because very low-frequency alleles are particularly sensitive to sequencing errors and distortions due to demographic processes or other model misspecifications. We inferred posterior distributions for each bootstrapped dataset, each one using the same 10^5^ sets of parameters randomly as prior to estimating summary statistics. We set the tolerance threshold in ABCreg to 0.025 such that 2.5 10^3^ values were accepted from posterior distributions.

### Forward-in-time simulations

We used SLiM 3 (Haller and Messer 2019) to generate simulated sequence variation data under our model and test the performances of our approach. We performed three different sets of simulations with and without the inclusion of non-equilibrium demographics, in which we modeled diverse rates of adaptation and BGS. For each set of simulations, we considered a 5.5 Myears branch length mimicking the human split from chimpanzee. In simulations accounting for demography, we added demographic events following a two-epoch model considering diverse expansion and bottleneck. We set the population size change to happen 1000 generations ago so testing *α*_(*x*)_ patterns depending on the SFS distortion due to recent demographic changes. We vary population size by a factor of 5 and 50 for a two-epoch expansion model and by 5 and 20 for a two-epoch bottleneck model. Note that unlike previous studies, we were interested in the *α*_(*x*)_ distortion under strong recent demographic changes (Eyre-Walker and Keightley 2009; Galtier 2016). In addition, we simulate a more *realistic* human demography following Tennessen *et al*. (2012) to model the variation in the 661 African individuals whose genomes are included in the 1000 Genome Project (Auton et al. 2015).

Each simulation represents a coding locus of 2 10^3^ bp flanked on each side by a 10^5^ bp loci, which contain deleterious alleles driving background selection. A total of 5 10^4^ genes were simulated, accounting for 10^8^*bp* of coding sequence. We performed the simulations following previously estimated values of negative selection of human proteins (Boyko et al. 2008), where the distribution of deleterious alleles follows a gamma-distribution with scale and shape parameters of 0.184 and 0.000402, respectively, which implies a mean fitness of 2*Ns* = −457 for negatively selected non-synonymous alleles. Strongly and weakly beneficial alleles followed a point-mass distribution given the population-scaled selection coefficients of 2*Ns* = 10 and 2*Ns* = 500, respectively. Our simulations supposed that 25% of mutations in each coding locus are synonymous while and 75% are non-synonymous. We used a mutation within each coding locus of *θ* = 4*Nµ* = 0.001 and a mean human recombination rate in the flanking sequence of *ρ* = 4*Nr* = 0.001. Given these parameters, we set an *α* value of 0.4 for equilibrium, and Tennessen simulations, and we decreased *α* to 0.2 for the two-epoch to vary the rate of adaptation tested. In the same way, because *α*_(*x*)_ pattern can also be affected by the number of individuals sampled (*n*), we tested different sample sizes for equilibrium (*n* = 500), Tennesen (*n* = 661) and two-epoch (*n* = 50) simulations. See Table 3 and Table 4 for parameter values of forward simulations. We rescaled ancestral population sizes (*N*_*e*_ = 10, 000) for equilibrium, and non-equilibrium model simulations according to our computational resources.

**Table 3.** SLiM simulated values to demographic equilibrium and Tennessen *et al*. (2012) simulations.

| Scenarios | $N_{anc}$ | $\mu_{flanking}$ | $r$ | $\mu_{coding}$ | $s_W^a$ | $s_S^a$ | $l_W^b$ | $l_S^c$ | $\alpha_W$ | $\alpha$ | $B$ |
| --- | --- | --- | --- | --- | --- | --- | --- | --- | --- | --- | --- |
| Demographic equilibrium | 500 | 1,32E-06 | 5,00E-07 | 5,00E-07 | 10 | 500 | 3,80E-03 | 3,90E-04 | 0,1 | 0,4 | 0,2 |
|  | 500 | 1,32E-06 | 5,00E-07 | 5,00E-07 | 10 | 500 | 7,70E-03 | 2,60E-04 | 0,2 | 0,4 | 0,2 |
|  | 500 | 1,32E-06 | 5,00E-07 | 5,00E-07 | 10 | 500 | 1,16E-02 | 1,30E-04 | 0,3 | 0,4 | 0,2 |
|  | 500 | 7,52E-07 | 5,00E-07 | 5,00E-07 | 10 | 500 | 3,80E-03 | 3,90E-04 | 0,1 | 0,4 | 0,4 |
|  | 500 | 7,52E-07 | 5,00E-07 | 5,00E-07 | 10 | 500 | 7,70E-03 | 2,60E-04 | 0,2 | 0,4 | 0,4 |
|  | 500 | 7,52E-07 | 5,00E-07 | 5,00E-07 | 10 | 500 | 1,16E-02 | 1,30E-04 | 0,3 | 0,4 | 0,4 |
|  | 500 | 1,83E-07 | 5,00E-07 | 5,00E-07 | 10 | 500 | 3,80E-03 | 3,90E-04 | 0,1 | 0,4 | 0,8 |
|  | 500 | 1,83E-07 | 5,00E-07 | 5,00E-07 | 10 | 500 | 7,70E-03 | 2,60E-04 | 0,2 | 0,4 | 0,8 |
|  | 500 | 1,83E-07 | 5,00E-07 | 5,00E-07 | 10 | 500 | 1,16E-02 | 1,30E-04 | 0,3 | 0,4 | 0,8 |
|  | 500 | 8,22E-10 | 5,00E-07 | 5,00E-07 | 10 | 500 | 3,80E-03 | 3,90E-04 | 0,1 | 0,4 | 0,999 |
|  | 500 | 8,22E-10 | 5,00E-07 | 5,00E-07 | 10 | 500 | 7,70E-03 | 2,60E-04 | 0,2 | 0,4 | 0,999 |
|  | 500 | 8,22E-10 | 5,00E-07 | 5,00E-07 | 10 | 500 | 1,16E-02 | 1,30E-04 | 0,3 | 0,4 | 0,999 |
| Tennesen model | 7310 | 1,32E-06 | 3,42E-08 | 3,42E-08 | 10 | 500 | 3,80E-03 | 3,90E-04 | 0,1 | 0,4 | 0,2 |
|  | 7310 | 1,32E-06 | 3,42E-08 | 3,42E-08 | 10 | 500 | 7,70E-03 | 2,60E-04 | 0,2 | 0,4 | 0,2 |
|  | 7310 | 1,32E-06 | 3,42E-08 | 3,42E-08 | 10 | 500 | 1,16E-02 | 1,30E-04 | 0,3 | 0,4 | 0,2 |
|  | 7310 | 7,52E-07 | 3,42E-08 | 3,42E-08 | 10 | 500 | 3,80E-03 | 3,90E-04 | 0,1 | 0,4 | 0,4 |
|  | 7310 | 7,52E-07 | 3,42E-08 | 3,42E-08 | 10 | 500 | 7,70E-03 | 2,60E-04 | 0,2 | 0,4 | 0,4 |
|  | 7310 | 7,52E-07 | 3,42E-08 | 3,42E-08 | 10 | 500 | 1,16E-02 | 1,30E-04 | 0,3 | 0,4 | 0,4 |
|  | 7310 | 1,83E-07 | 3,42E-08 | 3,42E-08 | 10 | 500 | 3,80E-03 | 3,90E-04 | 0,1 | 0,4 | 0,8 |
|  | 7310 | 1,83E-07 | 3,42E-08 | 3,42E-08 | 10 | 500 | 7,70E-03 | 2,60E-04 | 0,2 | 0,4 | 0,8 |
|  | 7310 | 1,83E-07 | 3,42E-08 | 3,42E-08 | 10 | 500 | 1,16E-02 | 1,30E-04 | 0,3 | 0,4 | 0,8 |
|  | 7310 | 8,22E-10 | 3,42E-08 | 3,42E-08 | 10 | 500 | 3,80E-03 | 3,90E-04 | 0,1 | 0,4 | 0,999 |
|  | 7310 | 8,22E-10 | 3,42E-08 | 3,42E-08 | 10 | 500 | 7,70E-03 | 2,60E-04 | 0,2 | 0,4 | 0,999 |
|  | 7310 | 8,22E-10 | 3,42E-08 | 3,42E-08 | 10 | 500 | 1,16E-02 | 1,30E-04 | 0,3 | 0,4 | 0,999 |

a population-scaled selection coefficient ( $2N_e s$ )
b Fixation probability of weakly beneficial alleles
c Fixation probability of strong beneficial alleles

**Table 4.** SLiM simulated values to the two-epoch model simulations.

| Scenarios | $N_{anc}$ | $\mu_{flanking}$ | $r$ | $\mu_{coding}$ | $s_W^a$ | $s_S^a$ | $l_W^b$ | $l_S^c$ | $\alpha_W$ | $\alpha$ | $B$ | | |
| --- | --- | --- | --- | --- | --- | --- | --- | --- | --- | --- | --- | --- | --- |
| Two-epoch model | 1000 | 50 | 1000 | 6,38E-08 | 2,50E-07 | 2,50E-07 | 10 | 500 | 2,91E-03 | 7,79E-05 | 0,1 | 0,2 | 0,4 |
|  | 1000 | 50 | 1000 | 6,38E-08 | 2,50E-07 | 2,50E-07 | 10 | 500 | 2,91E-03 | 7,79E-05 | 0,1 | 0,2 | 0,999 |
|  | 1000 | 200 | 1000 | 8,22E-10 | 2,50E-07 | 2,50E-07 | 10 | 500 | 2,91E-03 | 7,79E-05 | 0,1 | 0,2 | 0,4 |
|  | 1000 | 200 | 1000 | 6,38E-08 | 2,50E-07 | 2,50E-07 | 10 | 500 | 2,91E-03 | 7,79E-05 | 0,1 | 0,2 | 0,999 |
|  | 1000 | 5000 | 1000 | 8,22E-10 | 2,50E-07 | 2,50E-07 | 10 | 500 | 2,91E-03 | 7,79E-05 | 0,1 | 0,2 | 0,4 |
|  | 1000 | 5000 | 1000 | 8,22E-10 | 2,50E-07 | 2,50E-07 | 10 | 500 | 2,91E-03 | 7,79E-05 | 0,1 | 0,2 | 0,999 |
|  | 1000 | 50000 | 1000 | 6,38E-08 | 2,50E-07 | 2,50E-07 | 10 | 500 | 2,91E-03 | 7,79E-05 | 0,1 | 0,2 | 0,4 |
|  | 1000 | 50000 | 1000 | 6,38E-08 | 2,50E-07 | 2,50E-07 | 10 | 500 | 2,91E-03 | 7,79E-05 | 0,1 | 0,2 | 0,999 |

a population-scaled selection coefficient ( $2N_e s$ )
b Fixation probability of weakly beneficial alleles
c Fixation probability of strong beneficial alleles

### Human divergence and polymorphism

We also tested our model with empirical data. We retrieved polymorphic and divergence data in coding sequences from human hg38 assembly. Human assembly, coding sequences and annotations were retrieved from Ensembl release 109 (Cunningham et al. 2022). We estimated fixed substitutions in the human branch by Maximum Likelihood and ancestral sequence reconstruction based on human, chimpanzee, gorilla and orangutan alignments. First, we used pblat (Wang and Kong 2019) to align human CDS sequences on panTro6, gorGor6 and ponAbe3 assemblies downloaded from UCSC database (Nassar et al. 2023). To ensure that short exons at the edge of CDS were included, we used the pblat-fine option and a minimum identity threshold (-minIdentity) of 60%, keeping only the best scoring hit. The hit with the highest percentage of identity was retrieved if two or more equal best-score hits. Next, chimpanzee, gorilla and orangutan sequences were pblatted back to hg38 human assembly in order to make sure that the overall middle genomic position for each isoform was not greater than 1kb. By doing so, we finally retrieved 17,538 protein-coding genes accounting for a total of 94,030 CDS isoforms, which allowed us to identify the best reciprocal orthologous hits. Then, human, chimpanzee, gorilla and orangutan sequences were aligned using MACSE v2 to explicitly account for codon structure on protein-coding alignments (Ranwez et al. 2018). The total number of non-synonymous and synonymous fixations on the human branch were quantified from substitution maps using Hyphy v2.5 (Kosakovsky Pond et al. 2020). The process was divided into two steps: first, we fit the MG94xREV codon substitution model, a modification of the Muse and Gaut (1994) model, including a generalized time-reversible mutation rate matrix. The fit was done using MACSE alignments, and the great apes’ phylogenetic tree and phylogenies were re-estimated during the fitting allowing *d*_*N*_/*d*_*S*_ estimation by branch. Finally, each fitted model was used to reconstruct ancestral sequences and substitution maps. Fixations were summarized by gene taking into account all possible isoforms. Sites in multiple isoforms were counted only once, considering the most constraint annotation. Therefore, synonymous substitutions were considered non-synonymous if the same position were non-synonymous in at least one of the gene isoforms.

Polymorphic sites and derived allele frequencies were estimated from 1000GP phase 3 high-coverage phased data (Byrska-Bishop et al. 2022) across seven African ancestry populations. Non-synonymous and synonymous polymorphism were annotated using VEP (McLaren et al. 2016). Ancestral and derived allele frequencies were estimated using Ensembl v109 human ancestral allele information from EPO multi-alignments. Polymorphic sites only were counted if they overlapped those parts of human coding sequences that were previously aligned. In total, 17,538 orthologues were included in the analysis.

### Bootstrap test

We followed the bootstrap procedure extensively described at Enard and Petrov (2020) and Di *et al*. (2021) to compare adaptation levels between VIPs and non-VIP genes. The bootstrap procedure allows us to quantify adaptation specific to VIPs, comparing them with a control set that matches with VIPs on the number and average values of several confounding factors that also can determine the rate of adaptive evolution compared to the rest of the genome (Castellano et al. 2019b; Huang 2021). The bootstrap test relies on a straightforward control set-building algorithm that adds control genes to a progressively growing control set (Enard and Petrov 2020); in such a way, the growing control set has the same range of values of confounding factors as the tested set of genes of interest. Following this procedure, we avoid matching VIPs genes individually with non-VIPs control but match the overall VIPs set. Otherwise, we would drastically reduce the pool of potential control genes. When increasing the non-VIPs control set, we set a 5% matching limit over the VIPs set confounding average and limit each non-VIP to represent up to 1,2% of the control set. In total, we match nine potential confounding factors between VIPs and non-VIP control datasets (other confounding factors previously included in Di *et al*. (2021) were found to not impact the present comparison of VIPs and matched non-VIPs):

- Average overall expression in 53 GTEx v7 tissues (THE GTEX CONSORTIUM 2020). We used the log (in base 2) of TPM (Transcripts Per Million).
- Expression (log base 2 of TPM) in GTEx lymphocytes. Expression in immune tissues may impact the rate of sweeps.
- Expression (log base 2 of TPM) in GTEx testis. Expression in testis might also impact the rate of sweeps.
- deCode recombination rates in 50 kb and 500 kb gene-centered windows. The average recombination rates in the gene-centered windows are calculated using the most recent deCode recombination map (Halldorsson et al. 2019). We use both 50 kb and 500 kb window estimates to account for the effect of varying window sizes on the estimation of this confounding factor (same logic for other factors where we also use both 50 kb and 500 kb windows).
- GC content at the third position of codons in the aligned Ensembl coding sequences.
- The number of protein-protein interactions (PPIs) in the human protein interaction network (Luisi et al. 2015). The number of PPIs has been shown to influence the rate of sweeps (Luisi *et al*. 2015). We use the log (base 2) of the number of PPIs.
- The aligned coding sequence length.
- The proportion of immune genes. The matched control sets have the same proportion of immune genes as disease genes, immune genes being genes annotated with the Gene Ontology terms GO:0002376 (immune system process), GO:0006952 (defense response) and/or GO:0006955 (immune response) as of May 2020 (The Gene Ontology Consortium 2021).

## Results

### Evolutionary processes affecting adaptation inference

We used our forward simulations to retrieve polymorphism and divergence data with a priori known adaptation parameters. To input the data in our model, we pooled the SFS and number of fixations of 5 10^4^ genes. Note that this is about 2.5 times as many genes as appear in the human genome -we use this larger number of genes such that the trends in the simulated data will be clear and not dominated by noise, while noting that noise may play a larger role in real datasets for some species with limited proteome coverage.

As demonstrated previously, the frequency spectrum (Tataru et al. 2017) and *α*_(*x*)_ (Uricchio et al. 2019) are substantially affected by the presence of weakly beneficial alleles when the rate of such mutations is relatively high (see Figure 1, Figure 2). The asymptote of the *α*_(*x*)_ curve bends below the true value of *α*_(*x*)_, which is caused by an excess of high frequency non-synonymous variants under weak positive selection. This results in downward bias of *α* estimates when aMK-test (Messer and Petrov 2013) is applied to the data (see Table 5). In real sequencing datasets we cannot *a priori* separate beneficial and deleterious alleles, but in our simulated datasets we can remove the weakly beneficial alleles and test whether this will fix the downwards bias in aMK. When removing weakly beneficial alleles we observed an increase in *α* estimates from aMK which tend towards the true value of *α* (see Figure 1 an Table 5). In addition, *α*_(*x*)_ can be substantially affected by BGS, especially when weakly beneficial alleles contribute to the frequency spectrum (see Figure 1). In cases where *α* is dominated by strong adaptation, both asymptotic values (accounting for all alleles, or just neutral and deleterious) tend to be similar, because strongly beneficial alleles are not substantially impeded by selective interference with linked deleterious variation.

**Table 5.** *α* and aMK estimations.

| | True $\alpha_w$ | $B$ | True $\alpha$ | aMK<sub>(neutral + deleterious)</sub> | aMK<sub>(all alleles)</sub> |
| --- | --- | --- | --- | --- | --- |
| Demographic equilibrium | 0.1 | 0.2 | 0.194 | 0.163 | 0.132 |
|  | 0.2 | 0.2 | 0.178 | 0.161 | 0.102 |
|  | 0.3 | 0.2 | 0.161 | 0.151 | 0.058 |
|  | 0.1 | 0.4 | 0.273 | 0.222 | 0.185 |
|  | 0.2 | 0.4 | 0.253 | 0.214 | 0.141 |
|  | 0.3 | 0.4 | 0.238 | 0.191 | 0.078 |
|  | 0.1 | 0.8 | 0.369 | 0.337 | 0.294 |
|  | 0.2 | 0.8 | 0.352 | 0.317 | 0.222 |
|  | 0.3 | 0.8 | 0.335 | 0.308 | 0.172 |
|  | 0.1 | 0.999 | 0.401 | 0.358 | 0.312 |
|  | 0.2 | 0.999 | 0.386 | 0.338 | 0.241 |
|  | 0.3 | 0.999 | 0.370 | 0.354 | 0.203 |
| Tennesen model | 0.1 | 0.2 | 0.209 | 0.16 | 0.102 |
|  | 0.2 | 0.2 | 0.192 | 0.184 | 0.069 |
|  | 0.3 | 0.2 | 0.177 | 0.165 | 0.030 |
|  | 0.1 | 0.4 | 0.251 | 0.202 | 0.047 |
|  | 0.2 | 0.4 | 0.265 | 0.245 | 0.140 |
|  | 0.3 | 0.4 | 0.249 | 0.248 | 0.094 |
|  | 0.1 | 0.8 | 0.36 | 0.32 | 0.254 |
|  | 0.2 | 0.8 | 0.35 | 0.322 | 0.205 |
|  | 0.3 | 0.8 | 0.341 | 0.323 | 0.152 |
|  | 0.1 | 0.999 | 0.389 | 0.357 | 0.293 |
|  | 0.2 | 0.999 | 0.381 | 0.354 | 0.229 |
|  | 0.3 | 0.999 | 0.364 | 0.358 | 0.179 |

**Figure 1.**
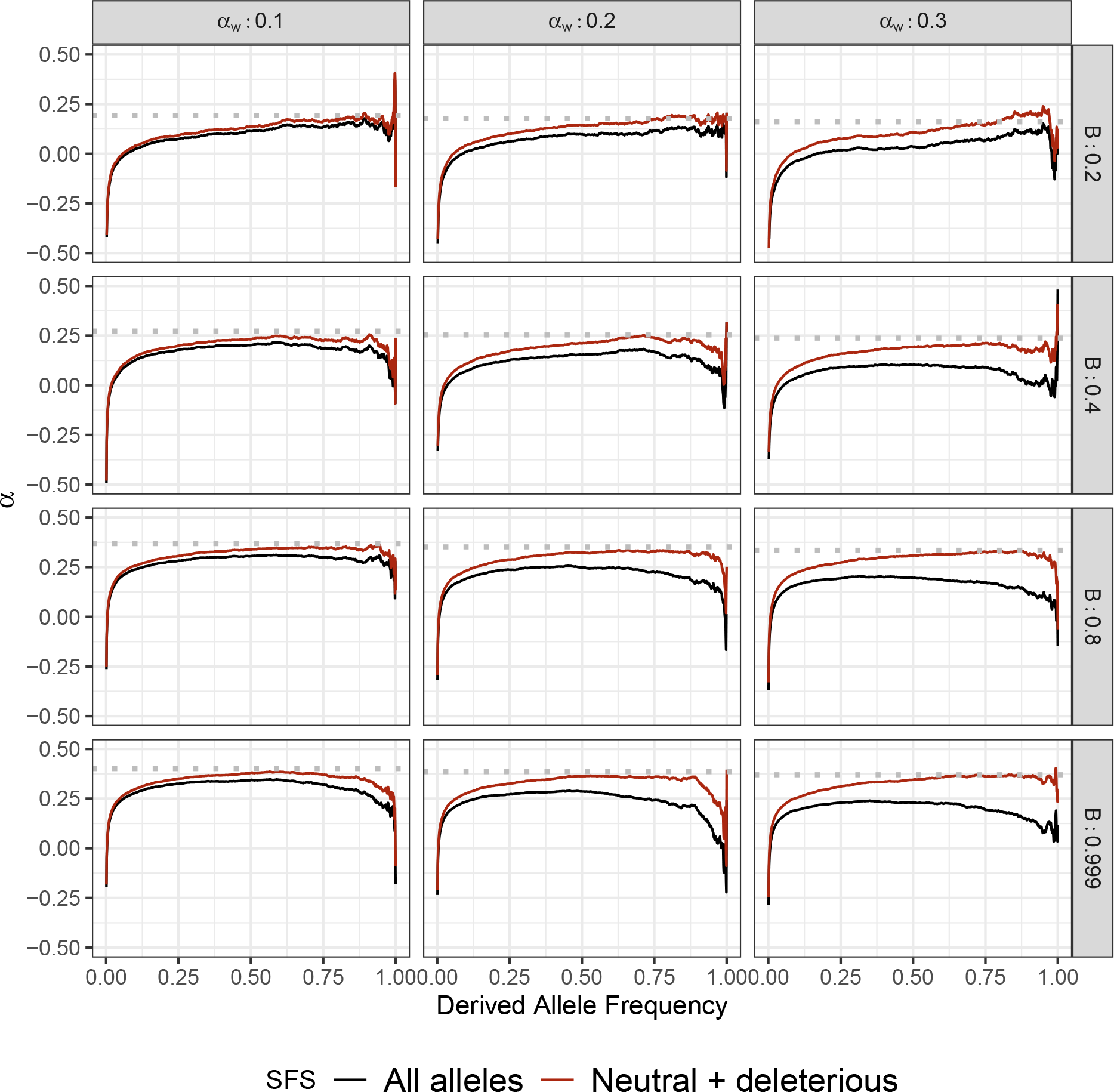
Effect of weakly advantageous mutation in the presence of BGS in equilibrium demography. We simulated the effect of weakly advantageous allele and BGS effect on *α*_(*x*)_. Each row represents a *B* value. Each column represents a proportion of *α*_*W*_ assuming a total proportion of adaptive substitutions of *α* = 0.4 in the absence of BGS. Dotted line is the true simulated *α*. Greater proportion of *α*_*W*_ largely biases *α*_(*x*)_ patterns which would affect aMK estimations. Stronger BGS removes beneficial mutations from *α*(*x*)

**Figure 2.**
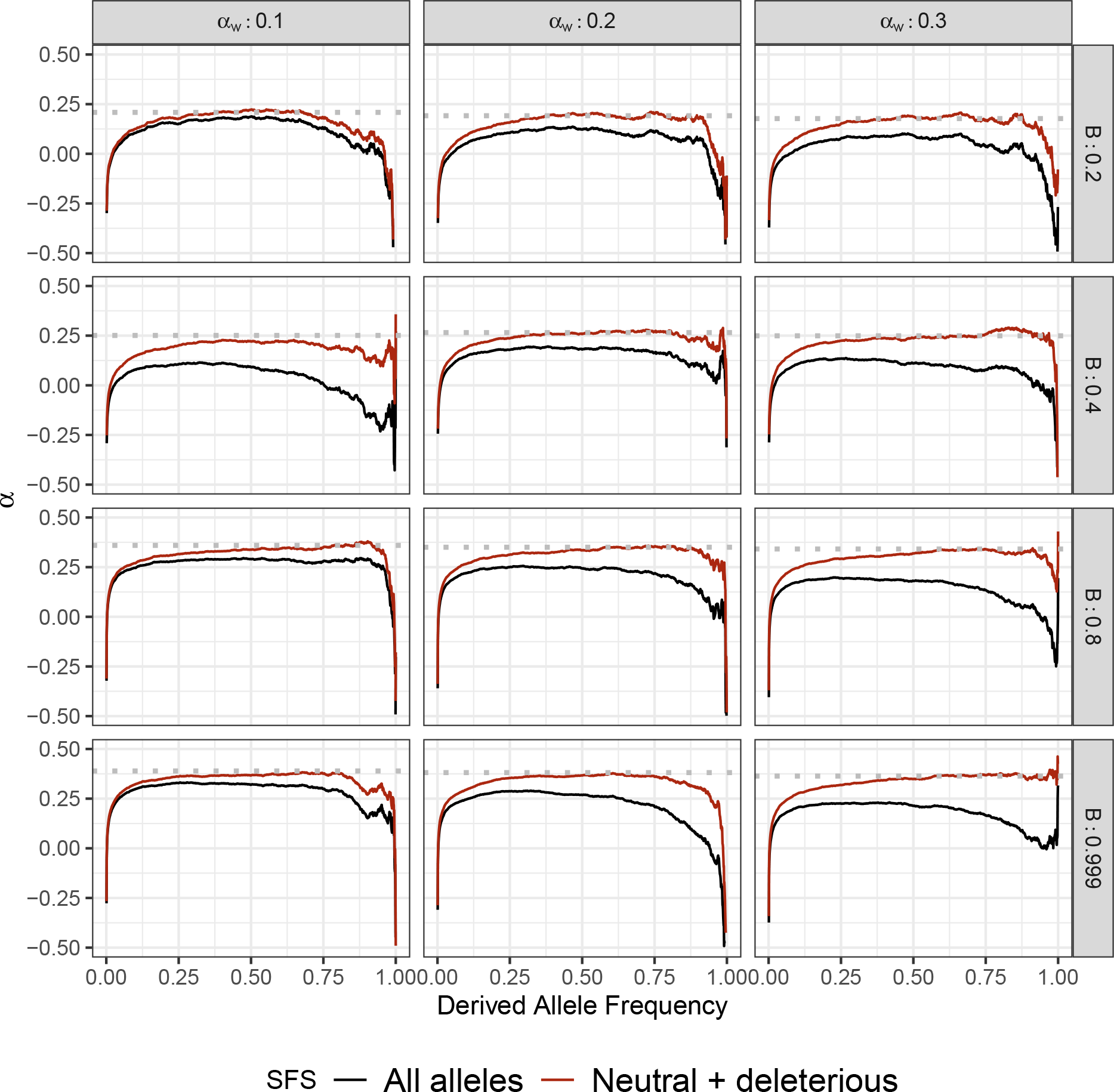
Effect of weakly advantageous mutation in the presence of BGS and under a non-equilibrium demography. We simulated the effect of weakly advantageous allele and BGS effect on *α*_(*x*)_ under the model of Tennessen *et al*. (2012). Each row represents a B value. Each column rep-resents a proportion of *α*W assuming a total proportion of adaptive substitutions of *α* = 0.4 in the absence of BGS. Dotted line is the true simulated *α*. Greater proportion of *α*_*W*_ largely biases *α*_(*x*)_ patterns which would affect aMK estimations. Stronger BGS removes beneficial mutations from *α*(*x*).

We also tested the effect of recent demographic events with a simulation of the Tennesen demographic model, specifically for the African continental group (Tennessen et al. 2012). We used demographic parameters following Adrion *et al*. (2020). To improve performance we simulated the African population in isolation, rather than including the full multi-population model –consequently our simulations do not include the effects of migration on the frequency spectrum. We observed similar patterns to equilibrium simulation regarding the overall shape and asymptotic values of *α*_(*x*)_ (Figure 2). Since this model includes a recent and rapid population expansion, we observe distortions to the frequency spectrum at extremely low and high frequencies due to the excess of rare alleles relative to an equilibrium demographic model.

The same effects were observed in the two-epoch expansion model. Overall we found the same expected differences on aMK and *α*_(*x*)_ curves in the two-epoch expansion considering *α* were reduced from 0.4 to 0.2 compared to equilibrium and Tennesen simulations (see Figure 3 and Table 6). Hence, BGS and the proportion of weakly advantageous polymorphism would drive the *α*_(*x*)_ curve differences despite the demographic model and the overall *α*. Nonetheless, as in Tennesen simulation, *α*_(*x*)_ are biased in extremely low and high frequencies due to the excess of rare alleles, depending on the event strength. However, like in the Tennesen simulation, aMK estimations are almost unaffected compared to the equilibrium simulation when removing weak advantageous mutations. That is not the case in the two-epoch bottleneck model. Although weakly advantageous polymorphism affects *α*_(*x*)_ curves similarly depending on the BGS strength, aMK estimations are biased compared to the equilibrium simulation. Since population contraction makes selection less efficient in removing deleterious mutations, such scenarios lead to an increment in *P*_*N*_ and underestimation of *α*, depending on the strength of the demographic event (see Figure 3 and Table 6). Because our method is based on the *α*_(*x*)_, such scenarios must be treated particularly carefully, and we discuss them in the following sections.

**Figure 3.**
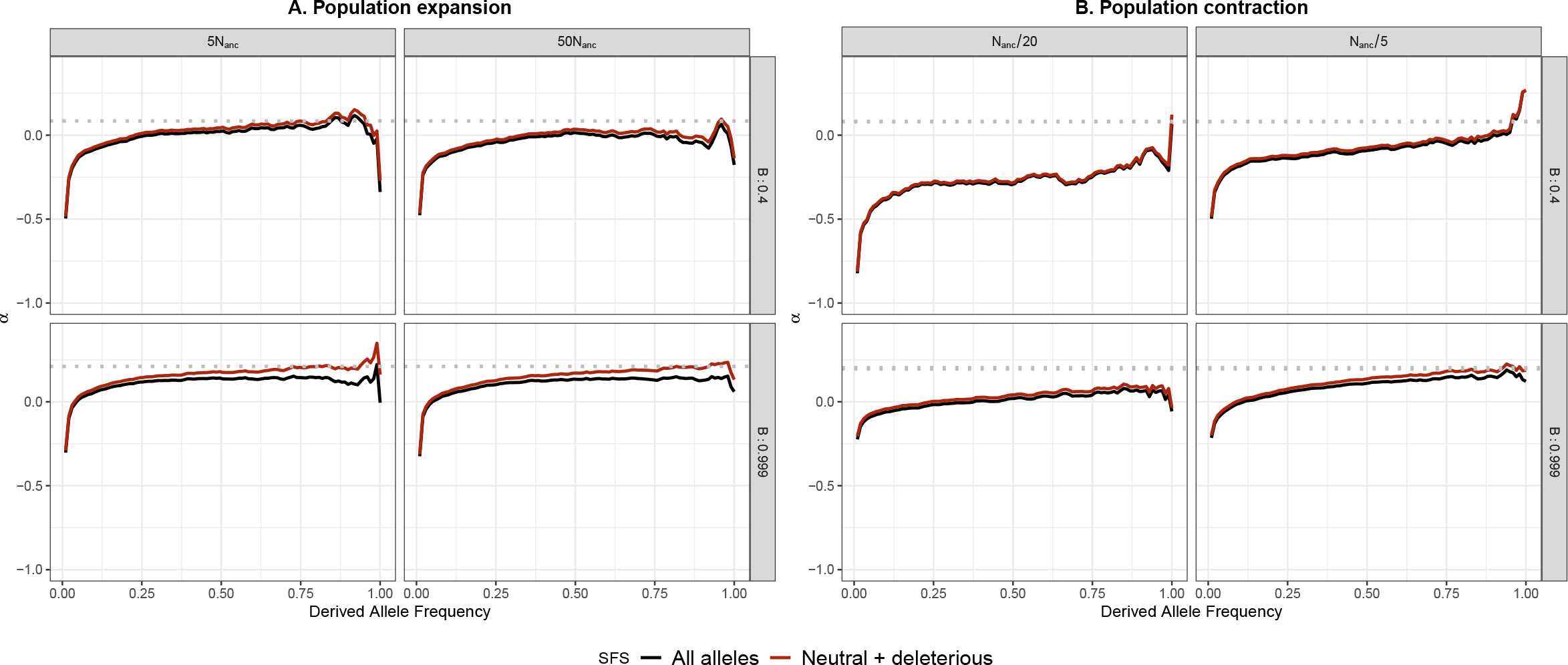
Effect of weakly advantageous mutation in the presence of BGS and population size change strength. We simulated the effect of weakly advantageous allele and BGS effect on *α*_(*x*)_ under a two-epoch model. Each row represents a B value. Each column represents the strength of the contraction or expansion of a proportion assuming a total proportion of adaptive substitutions of *α* = 0.2 in the absence of BGS. Dotted line is the true simulated *α. α*_*W*_ were set to contribute a half of the total *α*. Stronger BGS removes beneficial mutations from *α*(*x*).

**Table 6.** *α* and aMK estimations on the twoepoch model.

| | $N_{anc}$ | $N_B$ | True $\alpha_W$ | B | True $\alpha$ | aMK<sub>(neutral + deleterious)</sub> | aMK<sub>(all alleles)</sub> |
| --- | --- | --- | --- | --- | --- | --- | --- |
| Two-epoch<br>model | 1000 | 50.0 | 0.1 | 0.4 | 0.079 | -0.202 | -0.220 |
|  | 1000 | 50.0 | 0.1 | 0.999 | 0.195 | 0.077 | 0.050 |
|  | 1000 | 200.0 | 0.1 | 0.4 | 0.083 | 0.075 | 0.057 |
|  | 1000 | 200 | 0.1 | 0.999 | 0.204 | 0.188 | 0.147 |
|  | 1000 | 5000 | 0.1 | 0.4 | 0.084 | 0.059 | 0.030 |
|  | 1000 | 5000 | 0.1 | 0.999 | 0.210 | 0.194 | 0.130 |
|  | 1000 | 50000 | 0.1 | 0.4 | 0.085 | 0.013 | -0.014 |
|  | 1000 | 50000 | 0.1 | 0.999 | 0.212 | 0.189 | 0.129 |

### ABC evaluation using simulation

To compare true parameter values to inferred values, we estimated the mode and 95% CI for each posterior distribution that we obtained from ABC-MK as an approximation of *α*. Our method can distinguish both weak and strong adaptive contributions to *α* while performing reasonably accurate estimations. Table 7 shows inferred values and the associated error for each parameter and simulation. In all cases, the posterior distribution overlaps the distribution of the true values from bootstrapped datasets. Figure 6 presents the simulated values and mode estimates from posterior distributions on equilibrium simulations.

**Table 7.**
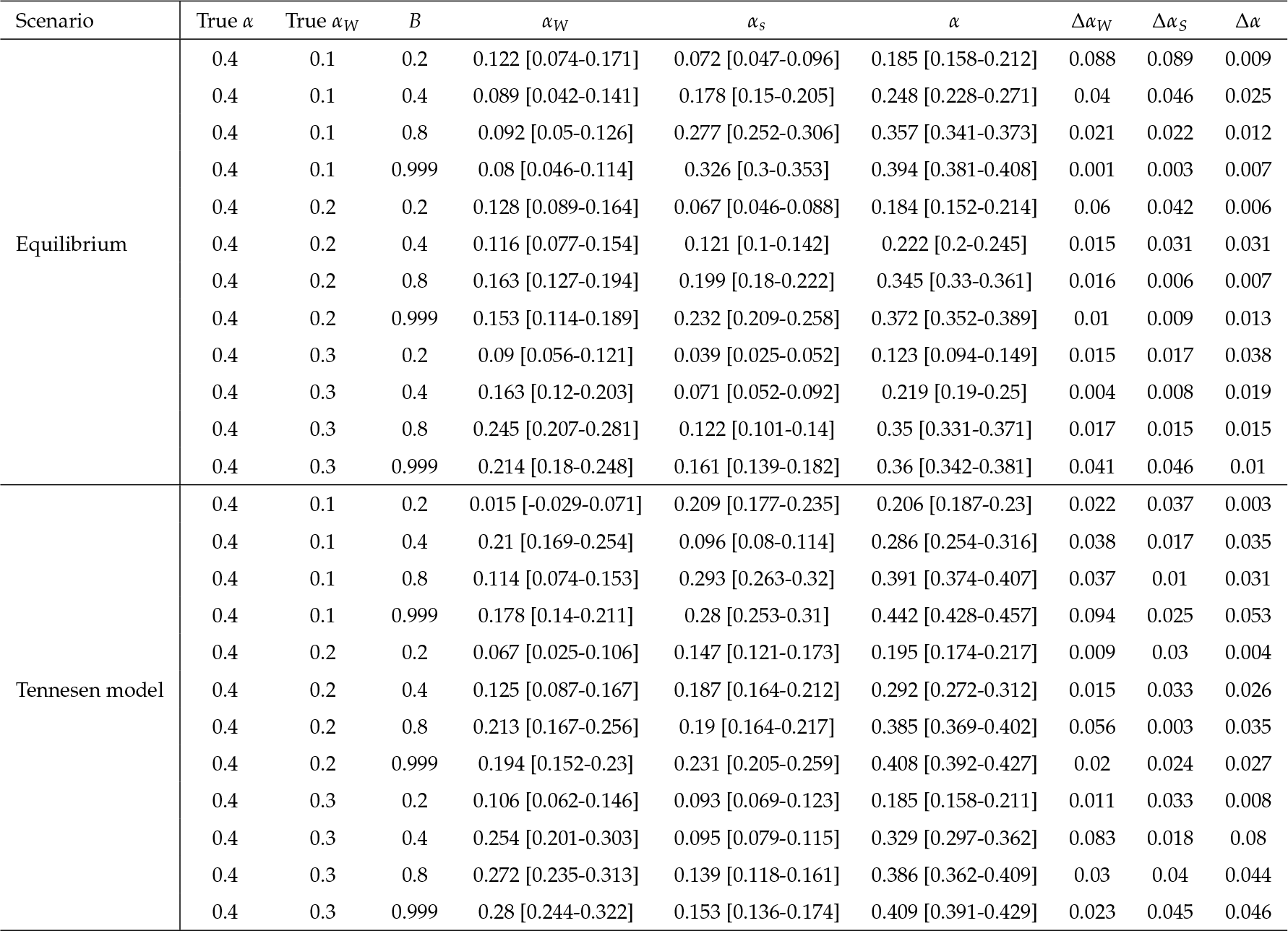
ABC inference in equilibrium and Tennessen *et al*. (2012) simulated datasets. Error were measured using the mean difference in 100 simulations replicas.

| Scenario | True $\alpha$ | True $\alpha_W$ | $B$ | $\alpha_W$ | $\alpha_S$ | $\alpha$ | $\Delta\alpha_W$ | $\Delta\alpha_S$ | $\Delta\alpha$ |
| --- | --- | --- | --- | --- | --- | --- | --- | --- | --- |
| Equilibrium | 0.4 | 0.1 | 0.2 | 0.122 [0.074-0.171] | 0.072 [0.047-0.096] | 0.185 [0.158-0.212] | 0.088 | 0.089 | 0.009 |
|  | 0.4 | 0.1 | 0.4 | 0.089 [0.042-0.141] | 0.178 [0.15-0.205] | 0.248 [0.228-0.271] | 0.04 | 0.046 | 0.025 |
|  | 0.4 | 0.1 | 0.8 | 0.092 [0.05-0.126] | 0.277 [0.252-0.306] | 0.357 [0.341-0.373] | 0.021 | 0.022 | 0.012 |
|  | 0.4 | 0.1 | 0.999 | 0.08 [0.046-0.114] | 0.326 [0.3-0.353] | 0.394 [0.381-0.408] | 0.001 | 0.003 | 0.007 |
|  | 0.4 | 0.2 | 0.2 | 0.128 [0.089-0.164] | 0.067 [0.046-0.088] | 0.184 [0.152-0.214] | 0.06 | 0.042 | 0.006 |
|  | 0.4 | 0.2 | 0.4 | 0.116 [0.077-0.154] | 0.121 [0.1-0.142] | 0.222 [0.2-0.245] | 0.015 | 0.031 | 0.031 |
|  | 0.4 | 0.2 | 0.8 | 0.163 [0.127-0.194] | 0.199 [0.18-0.222] | 0.345 [0.33-0.361] | 0.016 | 0.006 | 0.007 |
|  | 0.4 | 0.2 | 0.999 | 0.153 [0.114-0.189] | 0.232 [0.209-0.258] | 0.372 [0.352-0.389] | 0.01 | 0.009 | 0.013 |
|  | 0.4 | 0.3 | 0.2 | 0.09 [0.056-0.121] | 0.039 [0.025-0.052] | 0.123 [0.094-0.149] | 0.015 | 0.017 | 0.038 |
|  | 0.4 | 0.3 | 0.4 | 0.163 [0.12-0.203] | 0.071 [0.052-0.092] | 0.219 [0.19-0.25] | 0.004 | 0.008 | 0.019 |
|  | 0.4 | 0.3 | 0.8 | 0.245 [0.207-0.281] | 0.122 [0.101-0.14] | 0.35 [0.331-0.371] | 0.017 | 0.015 | 0.015 |
|  | 0.4 | 0.3 | 0.999 | 0.214 [0.18-0.248] | 0.161 [0.139-0.182] | 0.36 [0.342-0.381] | 0.041 | 0.046 | 0.01 |
| Tennesen model | 0.4 | 0.1 | 0.2 | 0.015 [-0.029-0.071] | 0.209 [0.177-0.235] | 0.206 [0.187-0.23] | 0.022 | 0.037 | 0.003 |
|  | 0.4 | 0.1 | 0.4 | 0.21 [0.169-0.254] | 0.096 [0.08-0.114] | 0.286 [0.254-0.316] | 0.038 | 0.017 | 0.035 |
|  | 0.4 | 0.1 | 0.8 | 0.114 [0.074-0.153] | 0.293 [0.263-0.32] | 0.391 [0.374-0.407] | 0.037 | 0.01 | 0.031 |
|  | 0.4 | 0.1 | 0.999 | 0.178 [0.14-0.211] | 0.28 [0.253-0.31] | 0.442 [0.428-0.457] | 0.094 | 0.025 | 0.053 |
|  | 0.4 | 0.2 | 0.2 | 0.067 [0.025-0.106] | 0.147 [0.121-0.173] | 0.195 [0.174-0.217] | 0.009 | 0.03 | 0.004 |
|  | 0.4 | 0.2 | 0.4 | 0.125 [0.087-0.167] | 0.187 [0.164-0.212] | 0.292 [0.272-0.312] | 0.015 | 0.033 | 0.026 |
|  | 0.4 | 0.2 | 0.8 | 0.213 [0.167-0.256] | 0.19 [0.164-0.217] | 0.385 [0.369-0.402] | 0.056 | 0.003 | 0.035 |
|  | 0.4 | 0.2 | 0.999 | 0.194 [0.152-0.23] | 0.231 [0.205-0.259] | 0.408 [0.392-0.427] | 0.02 | 0.024 | 0.027 |
|  | 0.4 | 0.3 | 0.2 | 0.106 [0.062-0.146] | 0.093 [0.069-0.123] | 0.185 [0.158-0.211] | 0.011 | 0.033 | 0.008 |
|  | 0.4 | 0.3 | 0.4 | 0.254 [0.201-0.303] | 0.095 [0.079-0.115] | 0.329 [0.297-0.362] | 0.083 | 0.018 | 0.08 |
|  | 0.4 | 0.3 | 0.8 | 0.272 [0.235-0.313] | 0.139 [0.118-0.161] | 0.386 [0.362-0.409] | 0.03 | 0.04 | 0.044 |
|  | 0.4 | 0.3 | 0.999 | 0.28 [0.244-0.322] | 0.153 [0.136-0.174] | 0.409 [0.391-0.429] | 0.023 | 0.045 | 0.046 |

We also tested our method using simulations performed under a two-epoch model, including moderate and strong ancient expansion and bottleneck demographic, and the demographic model of Tennessen *et al*. (2012) (see Table 7 and 8). Although the demographic events affect the number of segregating sites and the shape of the SFS (which is not modelled in our calculations), our method is reasonably robust to these distortions, including an ancient expansion in the ancestral population and a period of recent rapid growth when excluding low-frequency variants (DAC *<* 5 i.e. DAF *<* 0.0038; Figure 5, Figure 6, 7). The inference is biased towards weak selection when we include variants at which the SFS is most distorted, such as singletons and very high-frequency variants. However, the overall value of *α* is not strongly affected (Figure 4). Such a pattern likely reflects the qualitative similarity of recent growth events and weakly beneficial alleles in terms of their effects on the SFS, as both will disproportionately increase the number of non-synonymous variants at a low frequency relative to an equilibrium model. In Figure 4, we explore a more comprehensive range of parameters using the best-performing set of summary statistics (i.e., excluding derived allele counts under 5). Table 7 and Table 8 show the estimated parameters and their corresponding errors for both the Tennessen *et al*. (2012) and two-epoch model simulations.

**Table 8.** ABC inference in two-epoch model simulated datasets. Errors were measured using the mean difference in 100 simulations replicas.

| | $N_{anc}$ | $N_B$ | True<br>$\alpha$ | True<br>$\alpha_W$ | B | $\alpha_w$ | $\alpha_s$ | $\alpha$ | $\Delta\alpha_w$ | $\Delta\alpha_s$ | $\Delta\alpha$ |
| --- | --- | --- | --- | --- | --- | --- | --- | --- | --- | --- | --- |
| Two-epoch<br>model | 1000 | 50 | 0.2 | 0.1 | 0.4 | 0.04 [0-0.093] | 0.005 [-0.002-0.015] | 0.054 [0.01-0.12] | 0.007 | 0.045 | 0.03 |
|  | 1000 | 50 | 0.2 | 0.1 | 0.999 | -0.055 [-0.088-0.02] | 0.169 [0.137-0.205] | 0.107 [0.09-0.122] | 0.144 | 0.054 | 0.097 |
|  | 1000 | 200 | 0.2 | 0.1 | 0.4 | 0.106 [0.074-0.133] | 0.01 [0.006-0.013] | 0.123 [0.091-0.152] | 0.075 | 0.039 | 0.044 |
|  | 1000 | 200 | 0.2 | 0.1 | 0.999 | 0.005 [-0.009-0.021] | 0.029 [0.009-0.056] | 0.035 [0.013-0.058] | 0.082 | 0.08 | 0.16 |
|  | 1000 | 5000 | 0.2 | 0.1 | 0.4 | 0.05 [0.025-0.079] | 0.049 [0.026-0.077] | 0.1 [0.073-0.129] | 0.019 | 0.004 | 0.015 |
|  | 1000 | 5000 | 0.2 | 0.1 | 0.999 | 0.146 [0.094-0.197] | 0.117 [0.084-0.149] | 0.25 [0.232-0.272] | 0.055 | 0.002 | 0.04 |
|  | 1000 | 50000 | 0.2 | 0.1 | 0.4 | 0.005 [-0.011-0.025] | 0.028 [0.008-0.052] | 0.036 [0.014-0.066] | 0.026 | 0.026 | 0.049 |
|  | 1000 | 50000 | 0.2 | 0.1 | 0.999 | 0.058 [0.014-0.103] | 0.146 [0.108-0.182] | 0.195 [0.179-0.216] | 0.032 | 0.025 | 0.017 |

**Figure 4.**
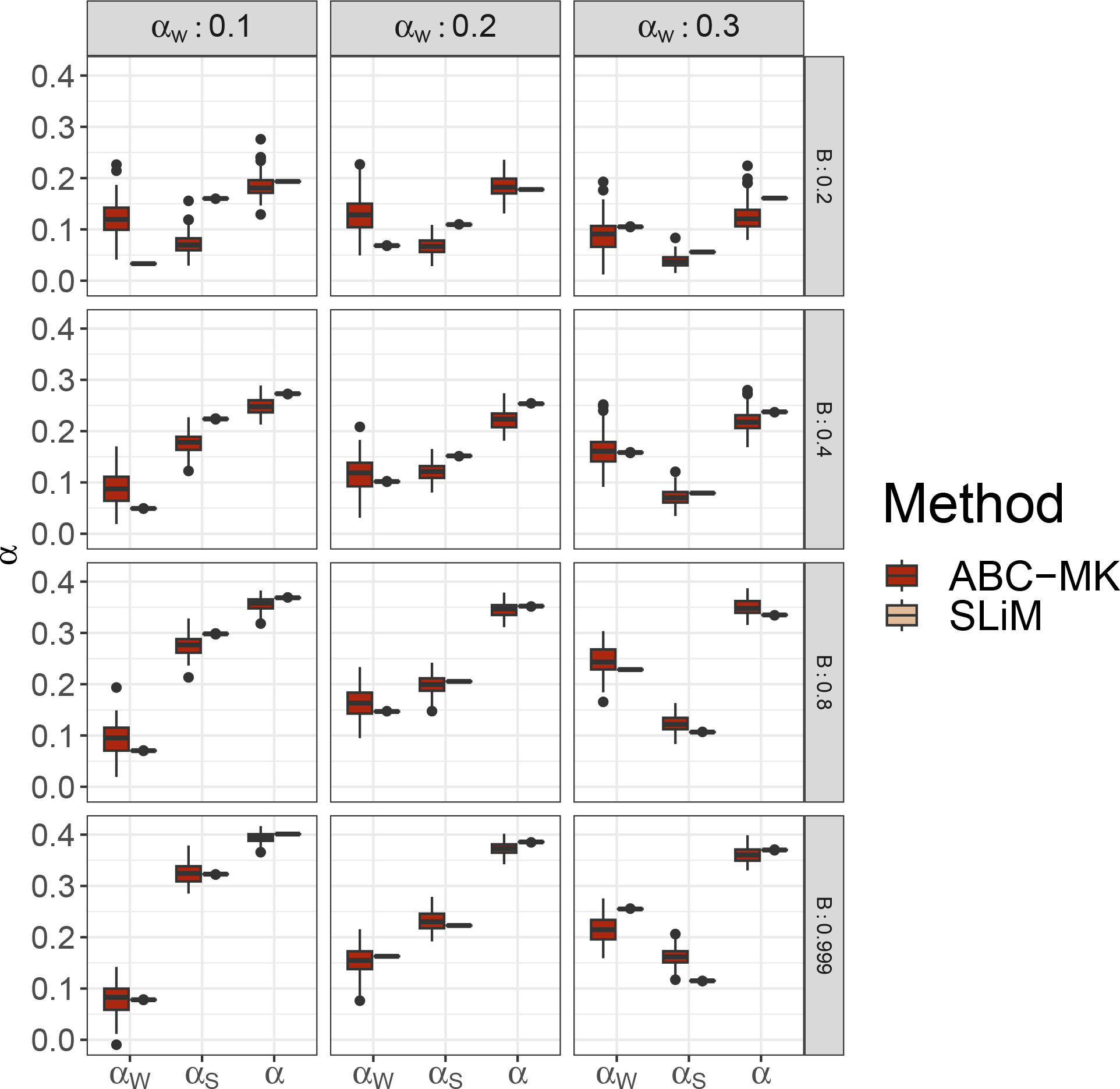
ABC-MK *α*_*W*_, *α*_*S*_ and *α* inferences at equilibrium. Mode distribution of 100 sets of summary statistics per parameter value. ABC inferences were performed using ABCreg Thornton (2009).

**Figure 5.**
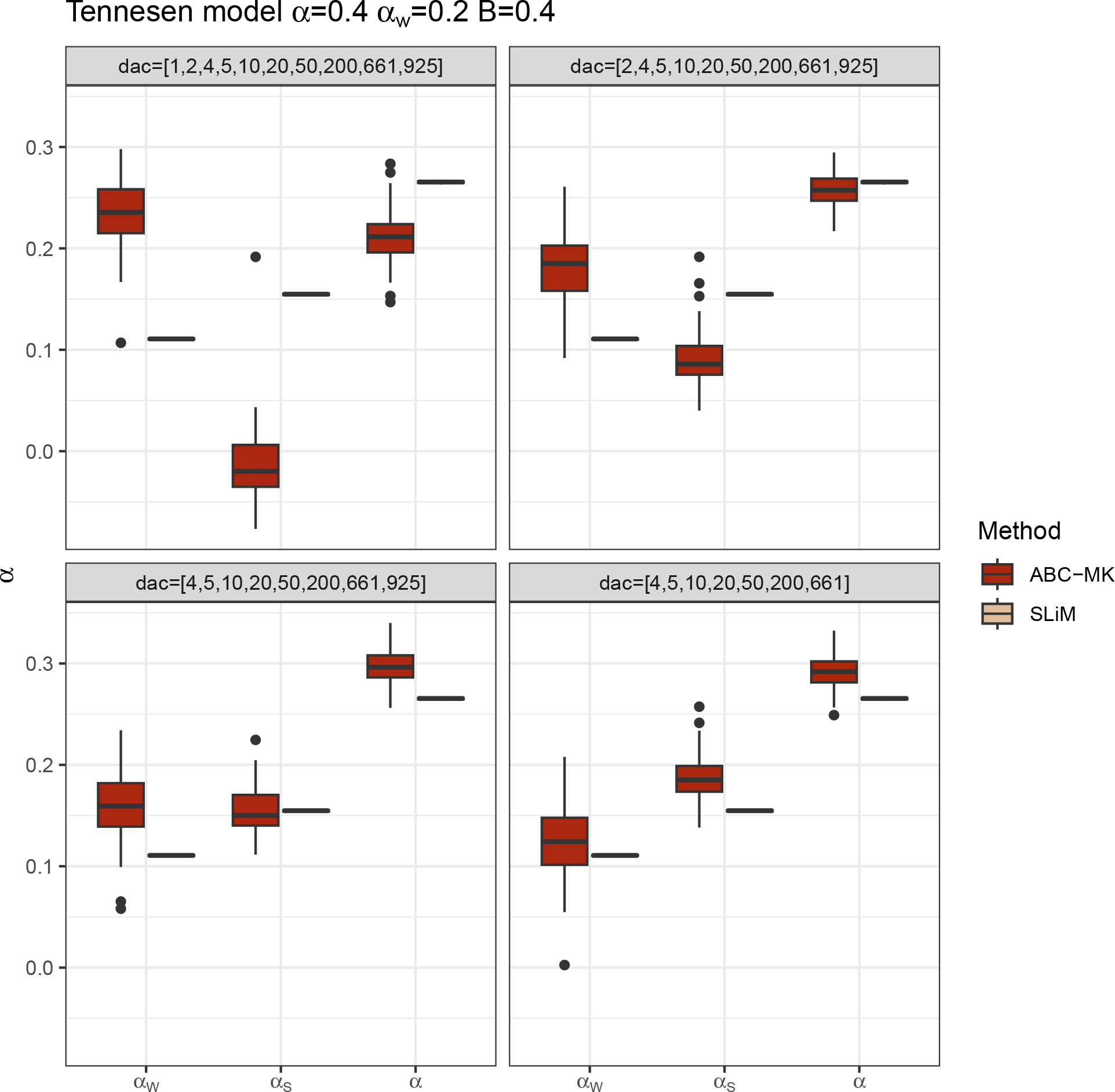
Summary statistics selection. ABC inference excluding low-frequency variants in Tennessen *et al*. (2012) demographic simulations.

**Figure 6.**
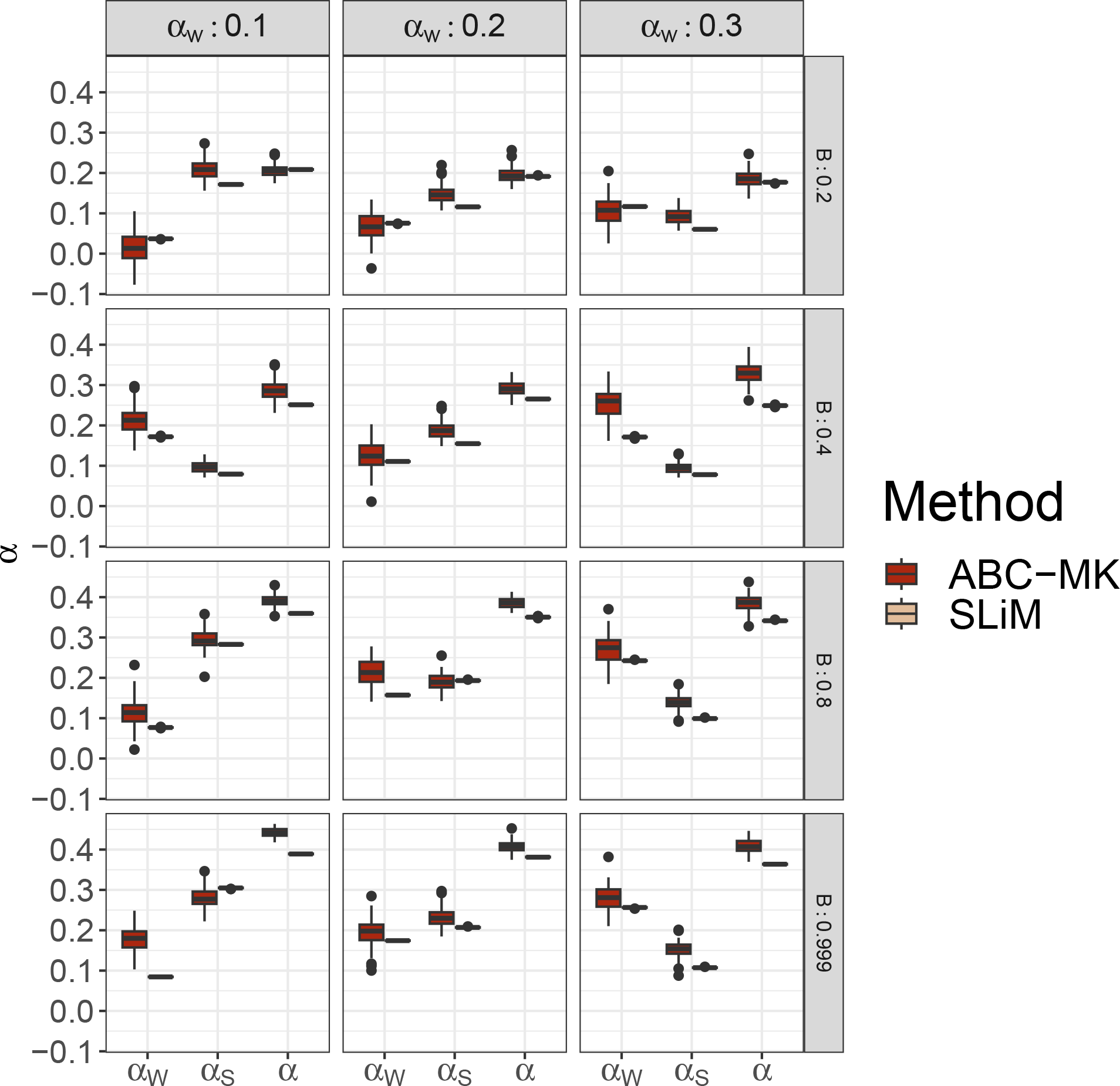
ABC-MK *α*_*W*_, *α*_*S*_ and *α* inferences of (Tennessen et al. 2012) demographic model simulations. Mode distribution of 100 datasets. ABC inference was performed using ABCreg Thornton (2009)

We noted that in cases where the strength of the bottleneck in the ancestral population is larger, our software performed worse *α* estimation than in the expansion scenarios (see Figure 7, Table 8). For such cases, we compared Grapes (Galtier 2016), a Maximum Likelihood software fitting the DFE to infer *α*, to our software. We run Grapes using GammaZero and GammaExpo models since both models best fit the simulated DFE (see Material and Methods). As shown in Figure 8, the GammaZero model and ABC-MK perform similarly in all cases, but the inference also fails when the ancient bottleneck and the BGS are strong. Besides, although the simulation included a higher proportion of weak adaptation, and the SFS included a significant amount of weakly adaptive variants (with parameters *α* = 0.2 and *α*_*W*_ = 0.1), we observed that the GammaExpo model provided less accurate estimations compared to the GammaZero model in all simulated scenarios (see Figure 7 and 3). Interestingly, although

**Figure 7.**
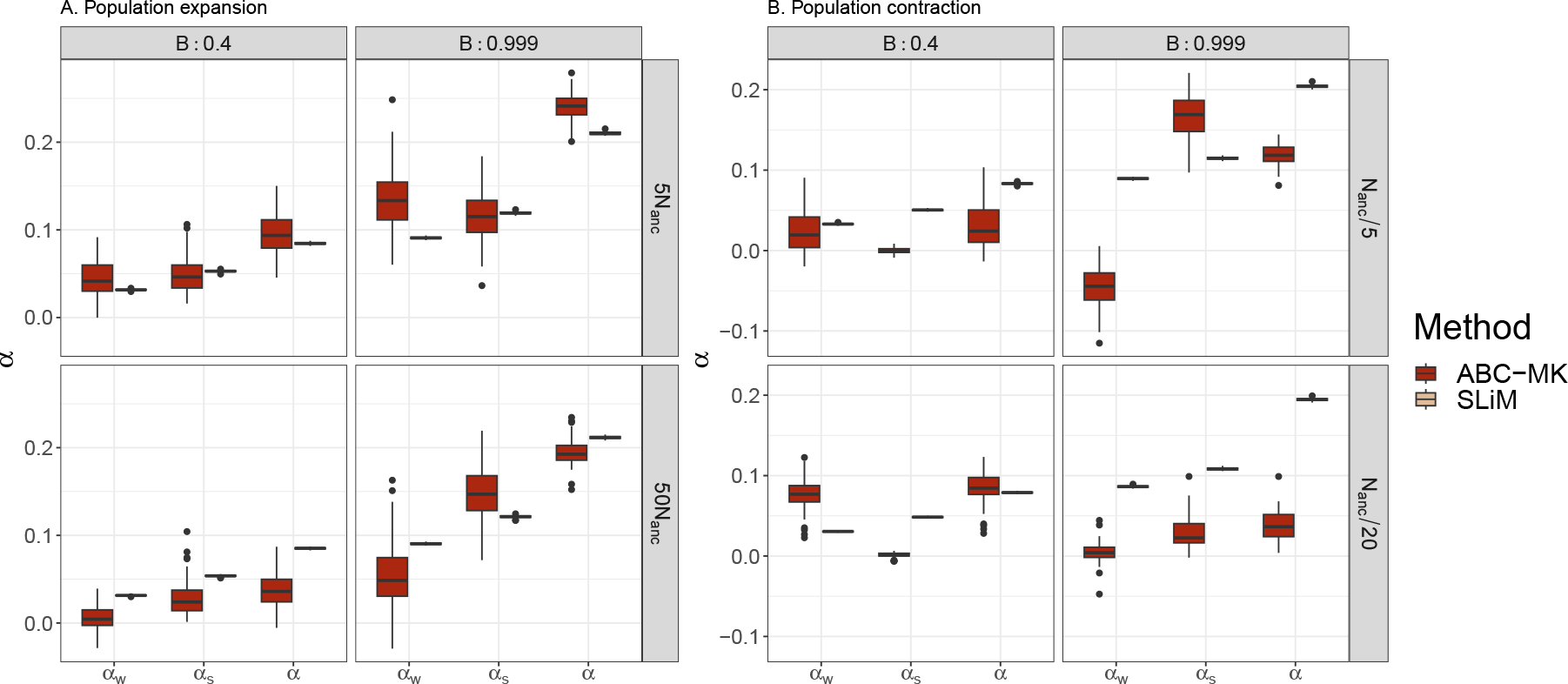
ABC-MK *α*_*W*_, *α*_*S*_ and *α* inferences of the two-epoch demographic model simulations. Mode distribution of 100 datasets. ABC inference was performed using ABCreg. A. Two-epoch expansion simulations. B. Two-epoch bottleneck simulations.

**Figure 8.**
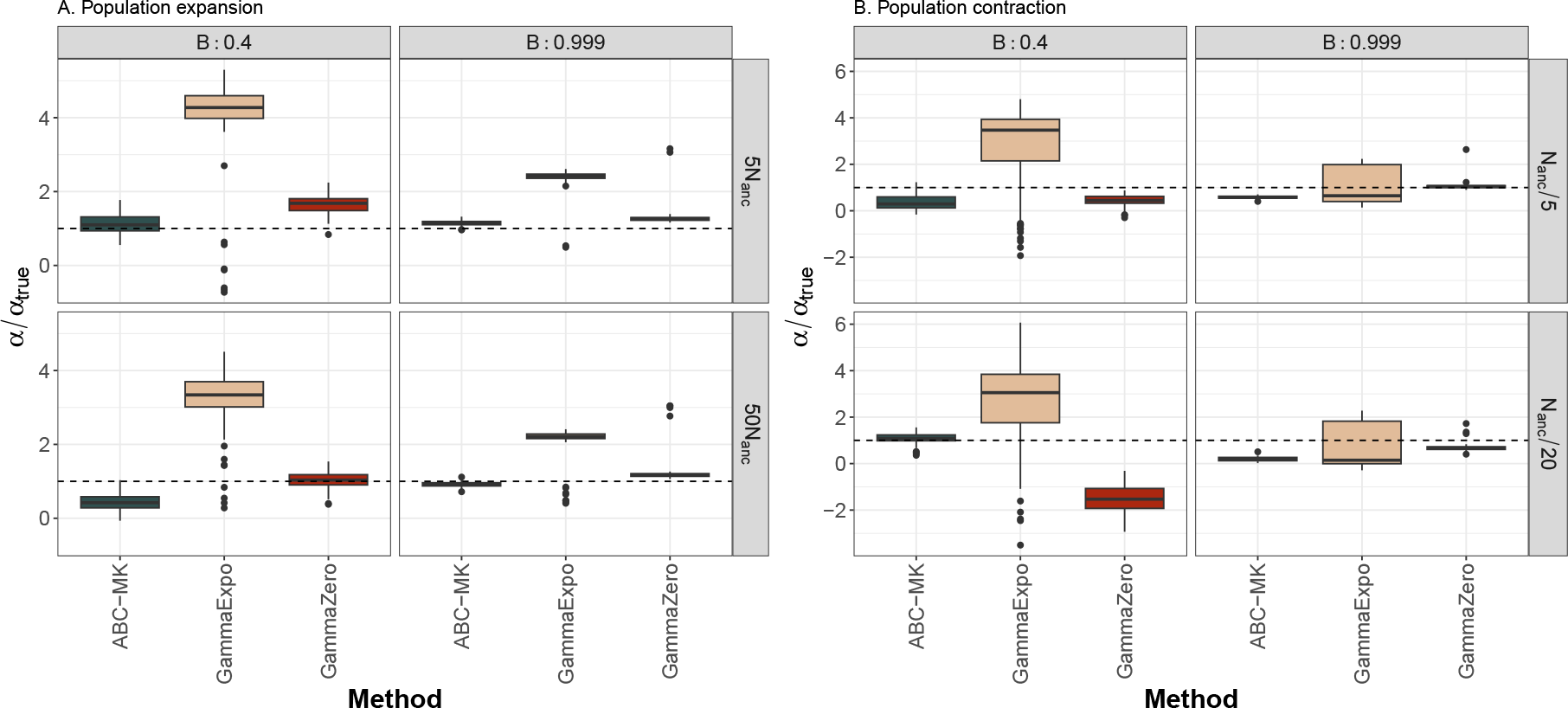
ABC-MK and Grapes *α* comparison of the two-epoch model simulations. A. Two-epoch expansion simulations. B. Two-epoch bottleneck simulations.

Grapes assumes independent segregating sites, we noted that it achieves accurate *α* estimations in the simulations, including BGS. Note that Grapes implement the *r* nuisance parameter correction proposed in Eyre-Walker *et al*. (2006). The *r* parameter was initially designed to capture the effect of demographic changes on the fate of synonymous and non-synonymous mutations (Eyre-Walker et al. 2006). In practice, the *r* parameter allows each SFS category to account for each own mutation rate (see equations 6 and 7 from Galtier (2016)). Considering the frequency category *i* at the SFS, *r*_*i*_ is estimated as the relative effective mutation rate of the *i* frequency compared to singletons at neutral sites (Eyre-Walker et al. 2006; Galtier 2016; Tataru et al. 2017). Multiple studies have shown that correcting the expected SFS by *r* can at least partially absorb other SFS distortions like the one produced by ascertainment bias, nonrandom sampling, population substructure or linked selection (Galtier 2016; Tataru et al. 2017; Galtier and Rousselle 2020). Similarly, we found that the *r* parameter can absorb unexpected polymorphic patterns in *α* estimates due to linked selection (Galtier 2016; Tataru et al. 2017; Galtier and Rousselle 2020). Still, as noted in Galtier and Rousselle (2020), we observed that absorbing linked selection distortions leads to a significant underestimation of the inferred *γ* parameter by Grapes, as well as DFE shape values, resulting in largely biased DFE model inferences in comparison with ABC-MK which recovers these parameters more precisely (see Figure 9, Figure 10, Table 9).

**Figure 9.**
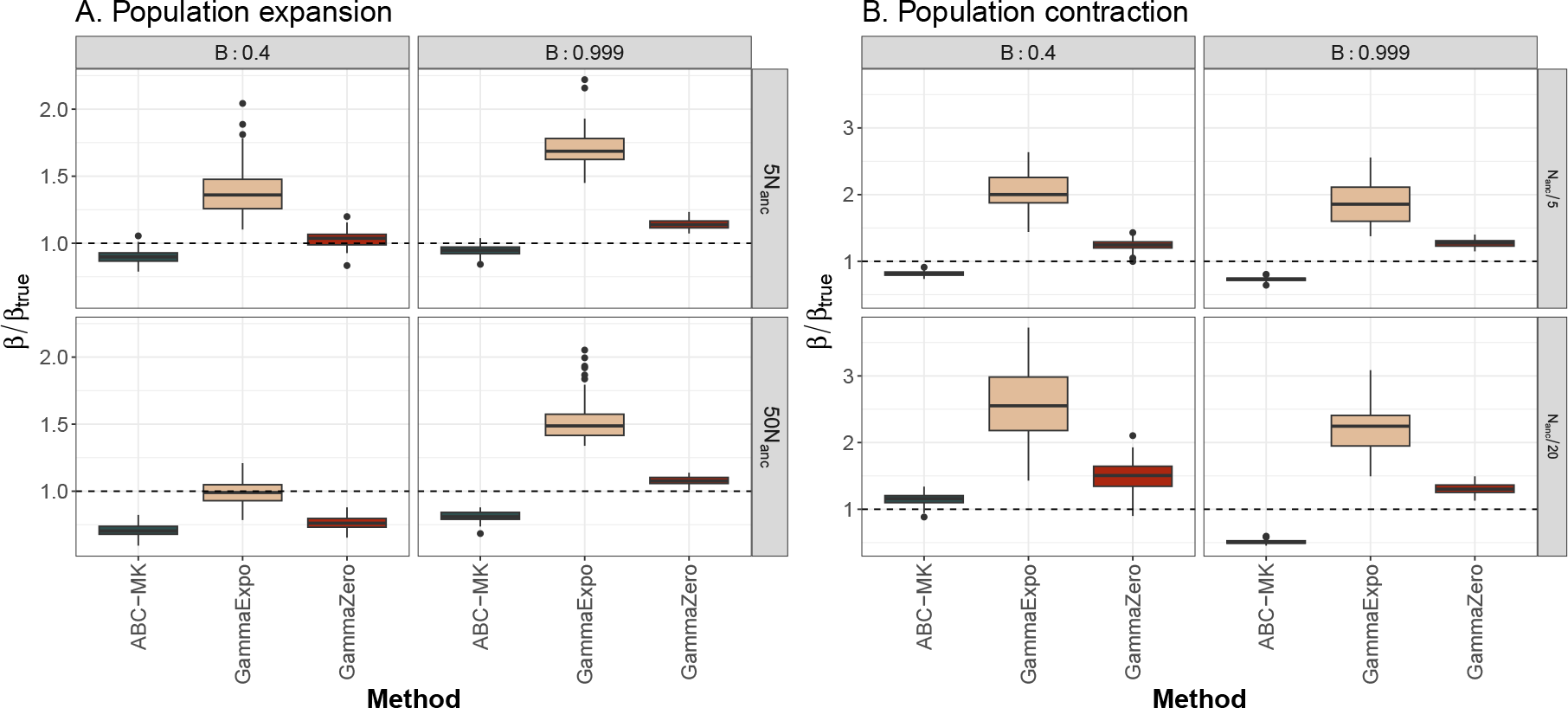
ABC-MK and Grapes comparison of the two-epoch model simulations. DFE shape (*β*) inference. A. Two-epoch expansion simulations. B. Two-epoch bottleneck simulations.

**Figure 10.**
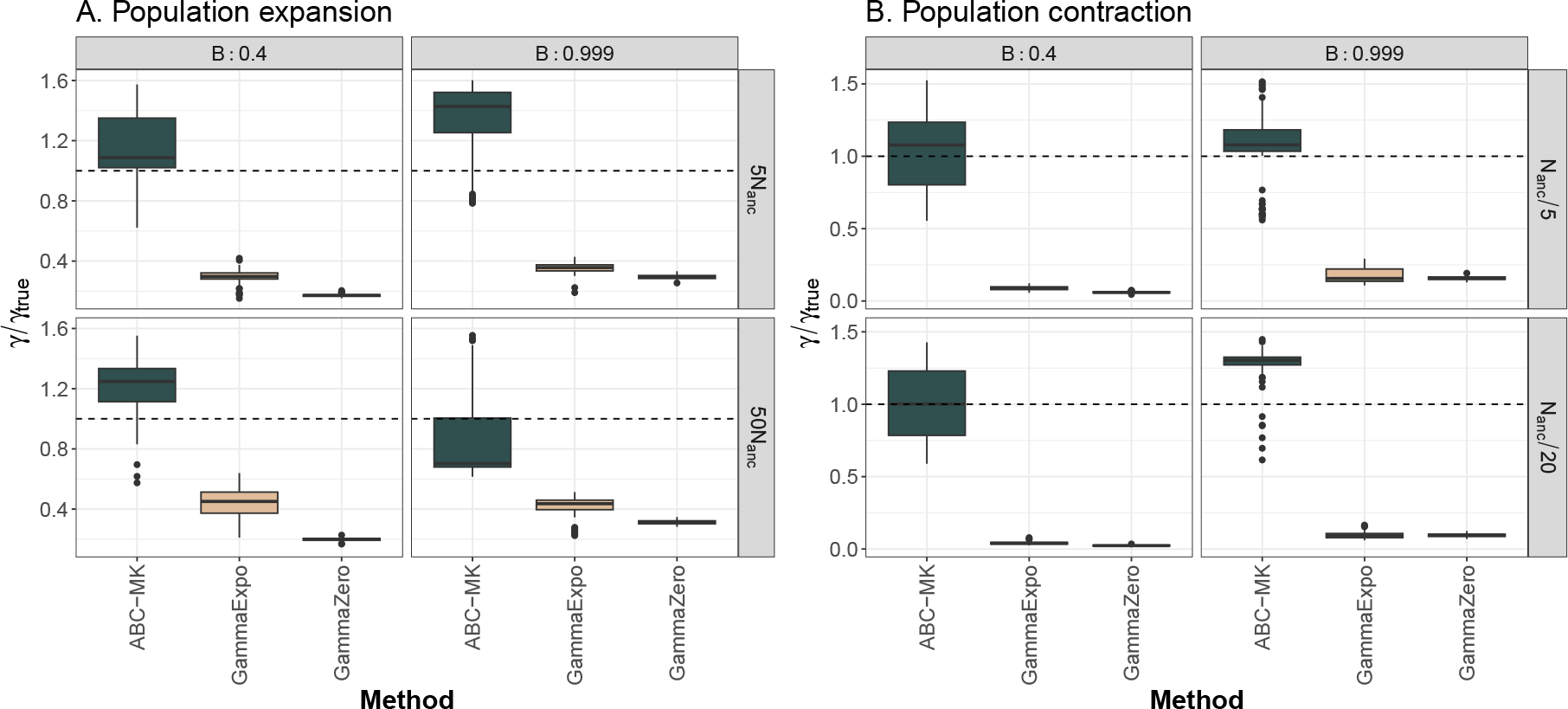
ABC-MK and Grapes comparison of the two-epoch model simulations. DFE negative selection coefficient (*γ*) inference. A. Two-epoch expansion simulations. B. Two-epoch bottleneck simulations.

**Table 9.** DFE inference in two-epoch model simulated datasets using ABC-MK and Grapes. Each method shows the mean ratio between th inference and the true value of 100 simulations replicas.

| Parameter | $N_{anc}$ | $N_B$ | True $\alpha$ | True $\alpha_W$ | B | True value | ABC-MK | GammaExpo | GammaZero |
| --- | --- | --- | --- | --- | --- | --- | --- | --- | --- |
| Shape ($\beta$) | 1000 | 50 | 0,2 | 0,1 | 0,4 | 0,184 | 0.827 | 2.048 | 1.242 |
|  | 1000 | 50 | 0,2 | 0,1 | 0,999 | 0,184 | 0.707 | 1.866 | 1.273 |
|  | 1000 | 200 | 0,2 | 0,1 | 0,4 | 0,184 | 1.126 | 2.556 | 1.509 |
|  | 1000 | 200 | 0,2 | 0,1 | 0,999 | 0,184 | 0.502 | 2.192 | 1.305 |
|  | 1000 | 5000 | 0,2 | 0,1 | 0,4 | 0,184 | 0.906 | 1.386 | 1.034 |
|  | 1000 | 5000 | 0,2 | 0,1 | 0,999 | 0,184 | 0.971 | 1.713 | 1.141 |
|  | 1000 | 50000 | 0,2 | 0,1 | 0,4 | 0,184 | 0.706 | 0.993 | 0.764 |
|  | 1000 | 50000 | 0,2 | 0,1 | 0,999 | 0,184 | 0.813 | 1.523 | 1.079 |
| Negative selection coefficient ($\gamma$) | 1000 | 50 | 0,2 | 0,1 | 0,4 | 457 | 1.253 | 0.089 | 0.060 |
|  | 1000 | 50 | 0,2 | 0,1 | 0,999 | 457 | 0.931 | 0.175 | 0.159 |
|  | 1000 | 200 | 0,2 | 0,1 | 0,4 | 457 | 1.103 | 0.041 | 0.024 |
|  | 1000 | 200 | 0,2 | 0,1 | 0,999 | 457 | 1.227 | 0.096 | 0.096 |
|  | 1000 | 5000 | 0,2 | 0,1 | 0,4 | 457 | 1.164 | 0.297 | 0.172 |
|  | 1000 | 5000 | 0,2 | 0,1 | 0,999 | 457 | 0.898 | 0.354 | 0.294 |
|  | 1000 | 50000 | 0,2 | 0,1 | 0,4 | 457 | 1.203 | 0.442 | 0.199 |
|  | 1000 | 50000 | 0,2 | 0,1 | 0,999 | 457 | 0.939 | 0.419 | 0.313 |

### Application of ABC-MK: human virus-interacting proteins

As an example application, we estimated adaptation in human genes that interact with viruses. Several studies have found that viral infections have driven adaptation in the human genome (e.g., Nédélec *et al*. (2016); Castellano *et al*. (2019b)). Genomic analysis of patterns of variation within experimentally determined Viral Interacting Proteins (VIPs) has repeatedly uncovered signals of both frequent and strong adaptation (Enard et al. 2016; Uricchio et al. 2019). Multiple lines of evidence suggest that the selective pressure imposed by viruses on hosts appears to be an important driver of strong adaptation during both long term (Enard et al. 2016; Castellano et al. 2019b; Uricchio et al. 2019) and more recent human evolution (Deschamps et al. 2016; Racimo et al. 2017; Enard and Petrov 2018).

We recently found that RNA viruses have driven more recent human evolution (in the past 50,000 years) at RNA virus-interacting VIPs (RNA VIPs) than DNA viruses at DNA virus-interacting VIPs (DNA VIPs) (Enard and Petrov 2018, 2020). Here we used ABC-MK to test whether the same was true during the millions of years of human evolution since divergence with chimpanzees. To this end, we limit our analyses to a subset of VIP and non-VIP mammals orthologs because gene age can significantly impact the rate of adaptation since they can evolve faster and experience mutations with stronger fitness effects to achieve their optimum fitness (Moutinho et al. 2022). Thus, to compare the impact of RNA and DNA viruses, we measured adaptation rates separately for the specific VIPs of nine different RNA viruses and the specific VIPs of six different DNA viruses each with more than 180 VIPs (Figure 11,12).

**Figure 11.**
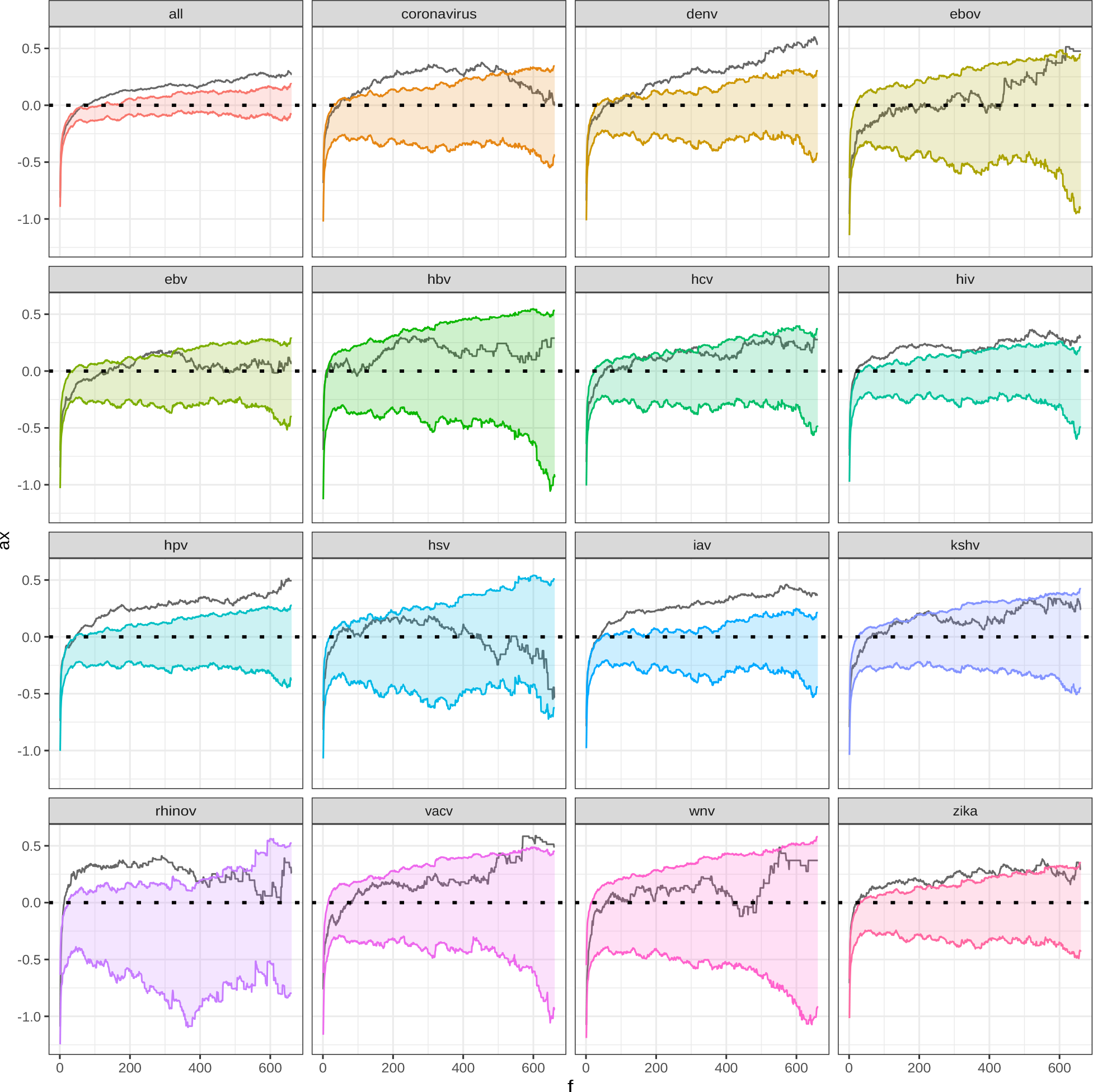
*α*_*x*_ patterns on VIPs and non-VIPs control. Colored range represent 95% *α*_(*x*)_ confidence interval.

**Figure 12.**
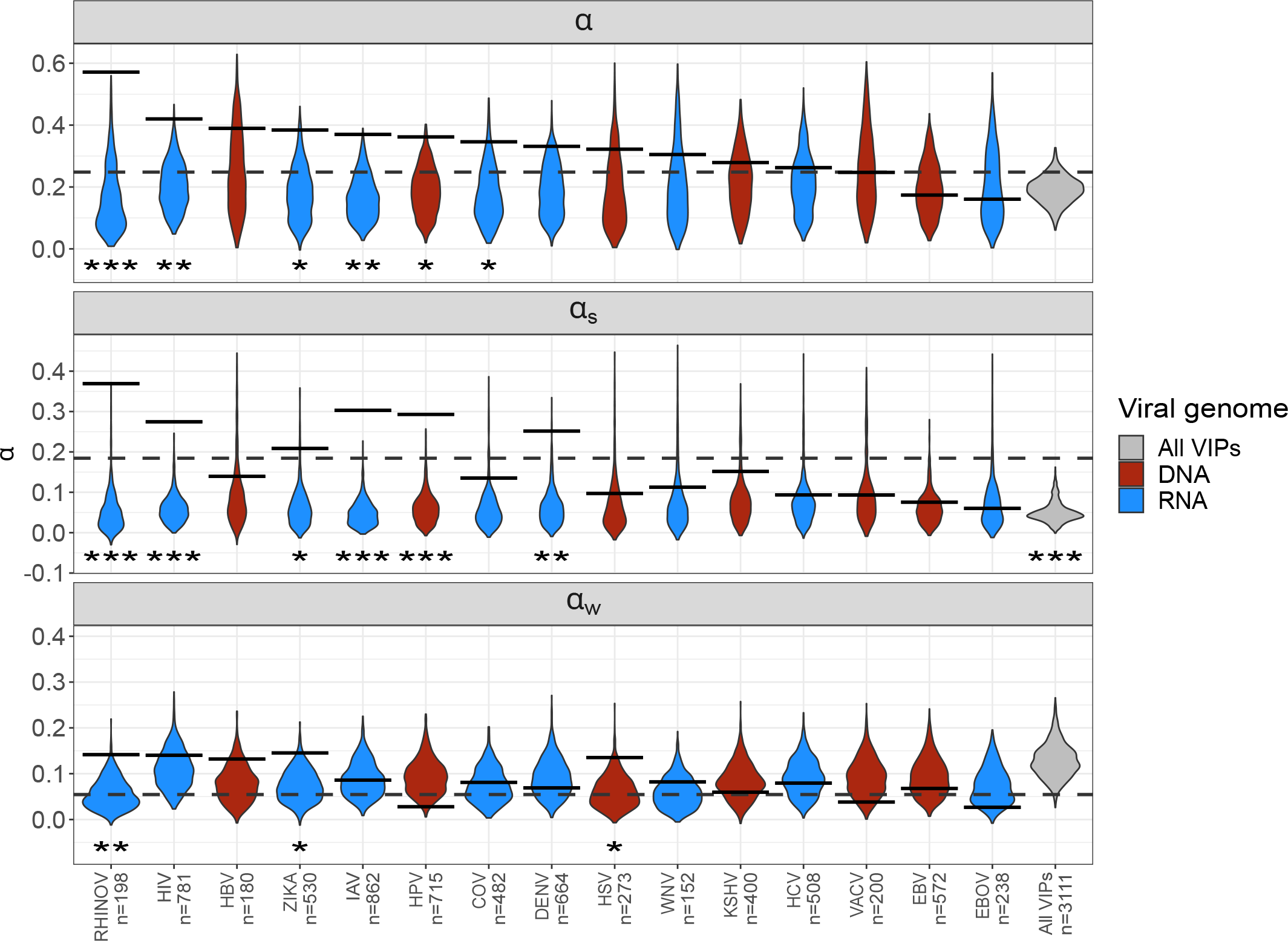
ABC-MK inference on different RNA and DNA-VIPs categories. Violin plots and solid lines represent inferences on non-VIP bootstrapped datasets and VIP categories according to the virus interaction. The dashed line is the overall VIPs inference.

We subset genomic data corresponding to the 661 individuals of African descent sampled in the Thousand Genomes Project, and fixed-substitution on human-branch were identified by maximum likelihood based on alignments of human (hg38 assembly), chimpanzee (panTro6 assembly) and orangutan (ponAbe3 assembly). Adaptation rates on VIPs were estimated following the previous bootstrap procedure described at Enard and Petrov (2020) and Di *et al*. (2021) (see Materials and Methods). The bootstrap procedure allows us to build random non-VIPs control sets that match the same average values of several confounding factors that might explain the adaptation level rather than the virus interactions themselves. We estimated non-VIPs null distributions by sampling 1,000 non-VIPs sets of the same size as the analyzed VIPs category. Because we defined we defined *P*_*N*(*x*)_ and *P*_*S*(*x*)_ as any polymorphisms above frequency *x*, we excluded any variants above 70% frequency to avoid mispolarization errors (see Figure 11). *α* values for VIPs and non-VIPs datasets were inferred through the mode and 95% CI estimates from posterior distributions following the previously described ABC scheme for each dataset.

We inferred values of *α, α*_*W*_, and *α*_*S*_ that were very similar to those inferred in Uricchio *et al*. (2019). Looking at the VIPs for specific RNA or DNA viruses, we find that a subset of RNA virus families drove the vast majority of adaptation in VIPs, (Figure 12). More specifically, RNA viruses drove strong adaptation but not weak adaptation (Figure 12), with five RNA viruses (RHINOV, HIV, ZIKAV, IAV and DENV) with VIPs with significantly elevated strong adaptation, versus only one DNA virus (HPV). We further find this is true using either *α* or *ω*_*a*_ (Figure 12, Table 10, Supplementary Figure 1, Supplementary Table 1) (Galtier 2016) to compare VIPs and controls. These results are coherent with growing evidence that RNA viruses drove strong recent adaptation more frequently than DNA viruses (Enard and Petrov 2018, 2020).

**Table 10.** Mode and 95% CI of *α*_*W*_, *α*_*S*_, *α*, negative selection coefficient (*γ* = 2*N*_*e*_*s*_−_), shape parameter (*β*) and p-values in human datasets.

| Dataset | Viral genome | $\alpha_W$ | $\alpha_S$ | $\alpha$ | $\gamma$ | $\beta$ | P-value $\alpha_W$ | P-value $\alpha_S$ | P-value $\alpha$ |
| --- | --- | --- | --- | --- | --- | --- | --- | --- | --- |
| RHINOV | RNA | 0.142 [0.033-0.664] | 0.369 [0.044-0.621] | 0.571 [0.36-0.841] | 471.161 [210.552-968.289] | 0.394 [0.299-0.504] | 0.001 | 0.001 | 0.009 |
| HIV | RNA | 0.14 [0.007-0.523] | 0.275 [-0.002-0.411] | 0.42 [0.285-0.623] | 567.58 [230.068-990.292] | 0.265 [0.213-0.358] | 0.006 | 0.001 | 0.237 |
| HBV | DNA | 0.132 [0.019-0.562] | 0.139 [0.006-0.475] | 0.389 [0.182-0.709] | 632.614 [231.317-993.048] | 0.244 [0.161-0.36] | 0.167 | 0.169 | 0.097 |
| ZIKA | RNA | 0.145 [0.011-0.517] | 0.209 [-0.012-0.405] | 0.384 [0.243-0.628] | 309.872 [223.871-982.288] | 0.252 [0.196-0.347] | 0.017 | 0.023 | 0.030 |
| IAV | RNA | 0.086 [-0.072-0.457] | 0.303 [0.001-0.44] | 0.37 [0.246-0.552] | 528.845 [215.823-977.793] | 0.263 [0.214-0.348] | 0.003 | 0.001 | 0.492 |
| HPV | DNA | 0.028 [-0.065-0.46] | 0.293 [-0.031-0.427] | 0.362 [0.217-0.527] | 404.733 [229.822-980.814] | 0.242 [0.182-0.322] | 0.015 | 0.001 | 0.981 |
| COV | RNA | 0.081 [-0.049-0.455] | 0.135 [0.002-0.411] | 0.346 [0.162-0.571] | 528.202 [219.173-988.197] | 0.242 [0.178-0.34] | 0.044 | 0.056 | 0.389 |
| DENV | RNA | 0.069 [-0.059-0.471] | 0.252 [-0.05-0.41] | 0.331 [0.188-0.523] | 724.572 [237.58-997.376] | 0.247 [0.185-0.333] | 0.054 | 0.007 | 0.670 |
| HSV | DNA | 0.135 [-0.003-0.507] | 0.097 [-0.011-0.379] | 0.322 [0.12-0.634] | 655.98 [223.942-976.381] | 0.236 [0.165-0.351] | 0.144 | 0.225 | 0.028 |
| WNV | RNA | 0.082 [-0.012-0.468] | 0.113 [-0.012-0.435] | 0.305 [0.089-0.613] | 322.908 [219.774-980.638] | 0.286 [0.209-0.404] | 0.195 | 0.173 | 0.240 |
| KSHV | DNA | 0.059 [-0.057-0.425] | 0.152 [-0.047-0.382] | 0.279 [0.103-0.518] | 484.308 [232.981-981.482] | 0.236 [0.172-0.335] | 0.266 | 0.089 | 0.727 |
| HCV | RNA | 0.079 [-0.029-0.42] | 0.094 [-0.037-0.332] | 0.262 [0.101-0.497] | 454.763 [213.725-974.311] | 0.219 [0.167-0.319] | 0.289 | 0.257 | 0.598 |
| VACV | DNA | 0.038 [-0.069-0.428] | 0.093 [-0.045-0.389] | 0.247 [0.036-0.519] | 536.851 [211.195-964.98] | 0.222 [0.147-0.331] | 0.476 | 0.364 | 0.917 |
| EBV | DNA | 0.068 [-0.032-0.368] | 0.076 [-0.027-0.266] | 0.174 [0.057-0.445] | 886.085 [226.868-980.951] | 0.21 [0.164-0.296] | 0.568 | 0.315 | 0.693 |
| EBOV | RNA | 0.027 [-0.052-0.383] | 0.06 [-0.043-0.331] | 0.16 [0.005-0.465] | 577.142 [227.886-994.858] | 0.241 [0.159-0.339] | 0.571 | 0.492 | 0.904 |
| All VIPs | DNA / RNA | 0.054 [-0.087-0.331] | 0.184 [-0.008-0.316] | 0.248 [0.165-0.414] | 339.151 [210.262-962.819] | 0.212 [0.182-0.281] | 0.093 | 0.001 | 0.988 |

## Data availability

We developed user-friendly software to run ABC-MK and the bootstrap procedure described at Enard and Petrov (2020) and Di *et al*. (2021). Software, tutorials and documentation are freely available at https://github.com/jmurga/MKtest.jl. The software is based on Julia language and supports multi-threading, interactive environments as well as Command Line Interface usage. Docker and Singularity containers are also available. The code and Jupyter notebooks used to perform the analyses are available at https://github.com/jmurga/abcmk_simulations.

## Funding

David Enard is funded by NIH NIGMS MIRA grant 5R35GM142677.

## Conflicts of interest

The author declares no conflict of interest.

## Supplementary material

**Supplementary Figure 1.**
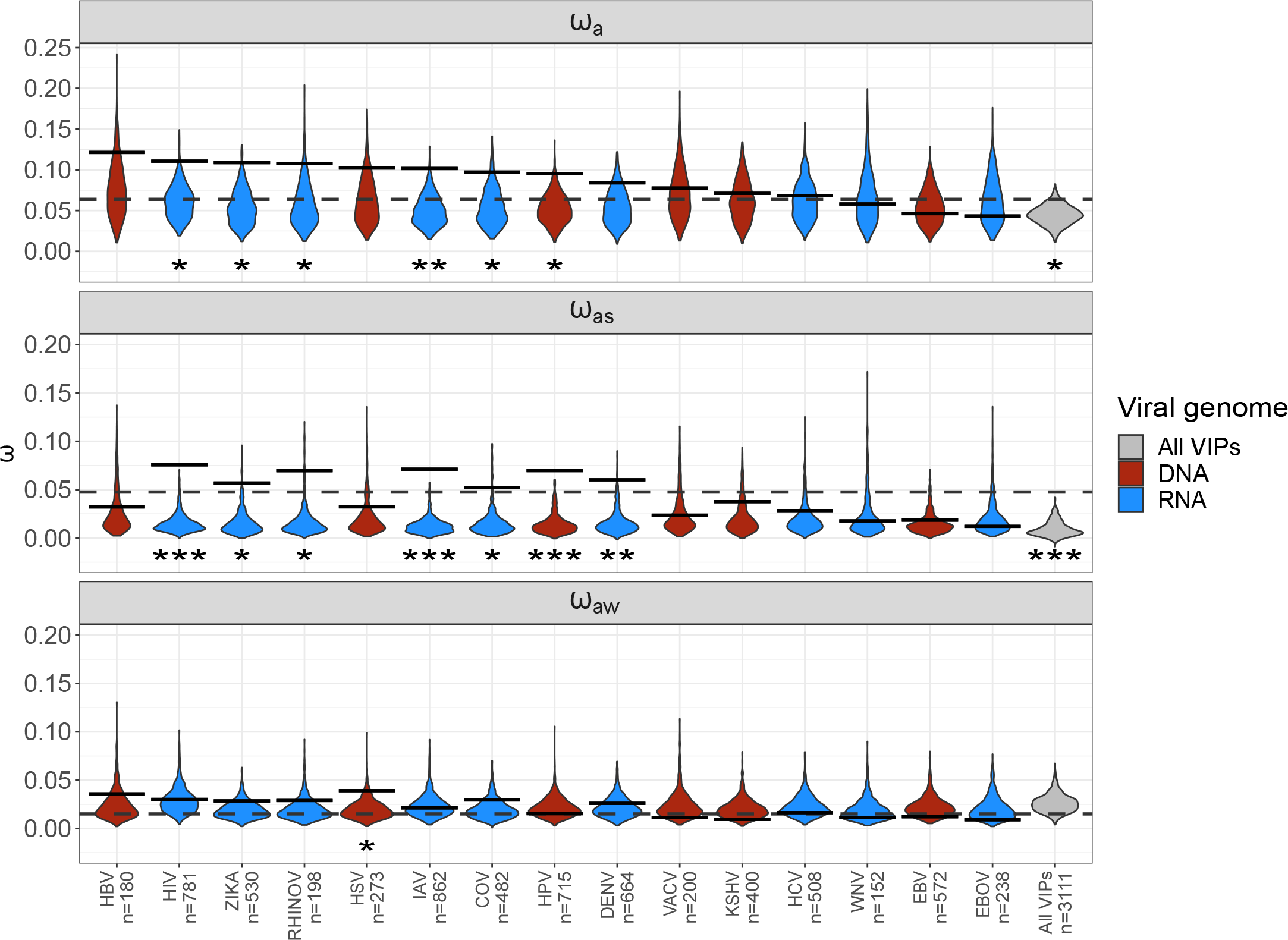
ABC-MK *ω*_*a*_ inference on different RNA and DNA-VIPs categories. Violin plots and solid lines represent inferences on non-VIP bootstrapped datasets and VIP categories according to the virus interaction. The dashed line is the overall VIPs inference.

**Supplementary Table 1.** Mode and 95% CI of *ω*_*aW*_, *ω*_*aS*_, *ω*_*a*_ and p-values in human datasets.

| Dataset | Viral genome | $\omega_{aW}$ | $\omega_{aS}$ | $\omega_a$ | P-value $\omega_{aW}$ | P-value $\omega_{aS}$ | P-value $\omega_a$ |
| --- | --- | --- | --- | --- | --- | --- | --- |
| RHINOV | RNA | 0.029 [-0.027-0.173] | 0.07 [-0.027-0.173] | 0.108 [0.029-0.219] | 0.049 | 0.018 | 0.161 |
| HIV | RNA | 0.03 [-0.01-0.159] | 0.076 [-0.015-0.139] | 0.111 [0.058-0.191] | 0.014 | 0.001 | 0.396 |
| HBV | DNA | 0.036 [-0.018-0.2] | 0.032 [-0.016-0.177] | 0.121 [0.023-0.257] | 0.092 | 0.265 | 0.176 |
| ZIKA | RNA | 0.029 [-0.01-0.168] | 0.057 [-0.016-0.144] | 0.109 [0.045-0.21] | 0.015 | 0.024 | 0.130 |
| IAV | RNA | 0.021 [-0.015-0.129] | 0.071 [-0.004-0.13] | 0.102 [0.058-0.164] | 0.005 | 0.001 | 0.496 |
| HPV | DNA | 0.015 [-0.013-0.122] | 0.07 [-0.007-0.134] | 0.096 [0.058-0.163] | 0.017 | 0.001 | 0.731 |
| COV | RNA | 0.03 [-0.021-0.146] | 0.052 [-0.009-0.146] | 0.097 [0.029-0.193] | 0.044 | 0.029 | 0.149 |
| DENV | RNA | 0.026 [-0.012-0.118] | 0.06 [-0.009-0.126] | 0.084 [0.045-0.161] | 0.102 | 0.010 | 0.296 |
| HSV | DNA | 0.039 [-0.022-0.184] | 0.032 [-0.021-0.146] | 0.102 [0.006-0.222] | 0.089 | 0.170 | 0.046 |
| WNV | RNA | 0.011 [-0.005-0.102] | 0.018 [-0.006-0.104] | 0.058 [0.006-0.145] | 0.537 | 0.437 | 0.778 |
| KSHV | DNA | 0.009 [-0.011-0.115] | 0.038 [-0.01-0.113] | 0.071 [0.014-0.158] | 0.341 | 0.147 | 0.909 |
| HCV | RNA | 0.016 [-0.006-0.122] | 0.028 [-0.011-0.101] | 0.068 [0.011-0.153] | 0.382 | 0.187 | 0.726 |
| VACV | DNA | 0.011 [-0.01-0.109] | 0.024 [-0.005-0.119] | 0.078 [0.01-0.16] | 0.367 | 0.356 | 0.882 |
| EBV | DNA | 0.012 [-0.007-0.107] | 0.019 [-0.008-0.08] | 0.046 [0.005-0.137] | 0.619 | 0.251 | 0.864 |
| EBOV | RNA | 0.009 [-0.009-0.095] | 0.012 [-0.007-0.086] | 0.043 [-0.0-0.129] | 0.706 | 0.661 | 0.888 |
| All VIPs | DNA / RNA | 0.015 [-0.014-0.088] | 0.048 [-0.004-0.093] | 0.064 [0.039-0.12] | 0.050 | 0.001 | 0.900 |

## Notes

### Competing Interest Statement

The authors have declared no competing interest.

### Summary of Updates

Tables and plots updated with proper CI.

